# Characterizing intra- and inter-tumor heterogeneity in Ovarian high-grade serous carcinoma subtypes using single-cell and spatial transcriptomics

**DOI:** 10.1101/2025.09.15.676244

**Authors:** Weishan Li, Laurie Grieshober, Jason Gertz, Adriana Ivich, Jennifer A. Doherty, Casey S. Greene, Stephanie C. Hicks

## Abstract

Ovarian high-grade serous carcinoma (HGSC) is an aggressive ovarian cancer with a heterogeneous tumor microenvironment (TME). Advances in single-cell RNA sequencing (scRNA-seq) and spatially-resolved transcriptomics have enabled the study of complex TME. This study explores connections between molecular subtypes described from bulk transcriptomes and spatial domains characterized by distinct gene expression in HGSC and their variability between patients. We explored both intra- and inter-tumor heterogeneity across 2D space and identified differing spatial patterns of gene expression pertaining to immune pathways and vasculature development. Functional characterization of tumor spaces revealed potentially shared cell states across molecular subtypes, while correlation analysis underscored subtype-specific spatial anti-colocalization between spots exhibiting antigen-presenting functions and B cell-mediated immunity. Lastly, we performed spatially-aware cell-cell communication analysis on the spatial samples and identified a molecular subtype specific difference in total signaling activity and heterogeneity in Midikine signaling between the differentiated subtype. Our results suggest that generating multiple tissue slices per patient might be necessary to enable comprehensive characterization of HGSC spatial transcriptomes.

## 1 Introduction

Ovarian high grade serous carcinoma (HGSC) is a common and highly aggressive type of ovarian cancer, which is generally associated with poor prognosis [1]. It is characterized by its loss of *TP53* and extensive copy number alterations [2]. HGSC also has a highly complex and heterogeneous tumor microenvironment (TME) broadly encompassing: 1) the parenchyma formed by epithelial cells and 2) the tumorassociated stroma from a collection of endothelial cells, inflammatory cells, cancer-associated fibroblasts, and stem/progenitor cells. HGSC tumor progression is a product of dynamic, reciprocal interaction between the parenchyma and growth supporting stroma [3].

Recent advances in single-cell RNA-sequencing (scRNA-seq) have shed new light on the HGSC TME [4, 5]. Molecular subtyping from bulk RNA-sequencing (RNA-seq) provides a framework for categorizing HGSC samples based on gene expression patterns that are associated with survival and may have therapeutic implications. Tothill et al. [6] was the first to report distinct molecular subtypes including the C1 subtype with strong stroma signature, the C2 subtype with elevated immune infiltration, the C3 subtype expressing a proliferative signature as well as the C5 subtype over-expressing mesenchymal genes. Subsequent studies expanded on this notion and built prediction algorithms to predict HGSC subtype based on gene expression microarrays [2, 7, 8]. The development of *consensusOV*, a R/Bioconductor package that implements four major methods for HGSC molecular subtype classification as well as a random forest-based consensus classifier, has facilitated generalizable and robust HGSC subtype prediction [9]. Another recent advancement was the creation of a platform-agnostic, RNA-seq compatible HGSC subtype clustering pipeline by [10], which also demonstrated consistent clustering of HGSC samples across ethnic groups.

More recently, molecular subtyping has been integrated into several HGSC gene expression atlases, enabling a deeper understanding of the role of stromal cell types and cell-cell interactions within each molecular subtype. For example, Olbrecht et al. [11] integrated *N* =7 HGSC scRNA-seq datasets and evaluated the contribution of the stromal cell type to the *k*=4 *consensusOV* molecular subtype signatures and identified differing cell-cell interactions. Deng et al. [12] applied *concensusOV* to *N* =5 HGSC scRNA-seq samples and reported subtyping results to be largely associated with cellular composition.

Furthermore, spatially-resolved transcriptomics offers a new perspective to understand the cellular composition in tumors by profiling gene expression *in situ*, which is lost in traditional scRNA-seq techniques [13]. This enables profiling tissue architecture of solid tumors and interactions between different cell types [14]. For example, Wu et al. [15] spatially profiled *N* =6 breast cancer samples and unraveled distinct cancer phenotypes, indicated by gene module enrichment scores, occupying mutually exclusive regions in space. Integrating scRNA-seq results with spatially resolved data can help address the resolution limitation of non-targeted capture and sequencing spatial transcriptomics platforms, such as 10x Genomics Visium [16].

Here, we investigated the intra-tumor heterogeneity observed in HGSC by spatially profiling four tissue sections from a single HGSC tumor, complemented by paired scRNA-seq data. We assessed the spatial organization of both biological pathways and key biomarkers. Next, we integrated our data with publicly available HGSC spatial data to explore differences in gene expression patterns and composition across known HGSC subtypes. We then investigated the spatial correlation of biological pathway and processes. We observed and characterized differences in the tumor architecture between tissue slices from the same tumor sample, as well as heterogeneous cell-cell interactions between tumor samples that, when analyzed by pseduobulking the gene expression counts (counts that have been normalized by library size and summed up across all spots in one sample), were attributed to the same transcriptomic subtype.

## 2 Results

### 2.1 Experimental design and overview of profiling one HGSC tumor sample

With a single fresh frozen HGSC tumor, we used the 10x Genomics Visium Spatial Gene Expression platform to spatially profile *N* =4 tissue sections from the same tumor to investigate the intra-tumor spatial heterogeneity (capture areas 1A, 2B, 3C, 4D). After applying QC metrics, the 1A capture area was removed from downstream analyses because there were less total UMI counts and detected genes per spot compared to other capture areas (**Figure S1**). In addition to the spatial profiling, we also leveraged a previously published paired scRNA-seq from the same tumor (16030X2) that was generated using the 10x Genomics Chromium platform [17]. After QC, the scRNA-seq dataset had 3469 cells for downstream analysis. Because this was a small number of cells, we integrated the paired scRNA-seq data with scRNA-seq data from an additional *N* =6 HGSC tumors (**Supplementary Table S1**) to ensure robust identification of cell types. All sequencing was performed at the Huntsman Cancer Institute (HCI) at the University of Utah. After this point, we refer to these samples as the Utah samples.

Five additional scRNA-seq samples published by Denisenko et. al [18] were retrieved from Gene Expression Omnibus (GEO) (GSE211956) and were subsequently integrated with the scRNA-seq samples generated by HCI to yield a dataset for use as cell type reference for deconvolution.

### 2.2 Characterizing intra-tumor heterogeneity using spatial transcriptomics

Our first objective was to characterize the spatial gene expression landscape across tissue slices derived from the same tumor tissue and assess intra-tumor heterogeneity (**Figure 1**). We investigated this at the spot-level and at the gene-level. Towards assessing the heterogeneity at the spot-level, we used a variable Bayes Non-negative Matrix Factorization (NMF) algorithm from the R/Bioconductor package *ccfindR* [19] to classify spots into *k*=10 spatial domains (**Figure 1A-C**). All 3 samples had around 2000 in-tissue spots, yet had strikingly different compositions of predicted spatial domains, despite originating from the same tumor tissue (**Figure 1D**). In addition, we identified spatially variable genes using Moran’s I to quantitatively demonstrate the intra-tumor heterogeneity between the capture areas. Similar to the differences in composition across the capture areas, we observe stronger association between Moran’s I values between capture area 3C and 4D, while capture area 2B exhibited heightened spatial-autocorrelation of immunoglobulin genes (*IGKC, IGHG3, IGLC1*) relative to both 3C and 4D (**Figure S2**).

**Figure 1:**
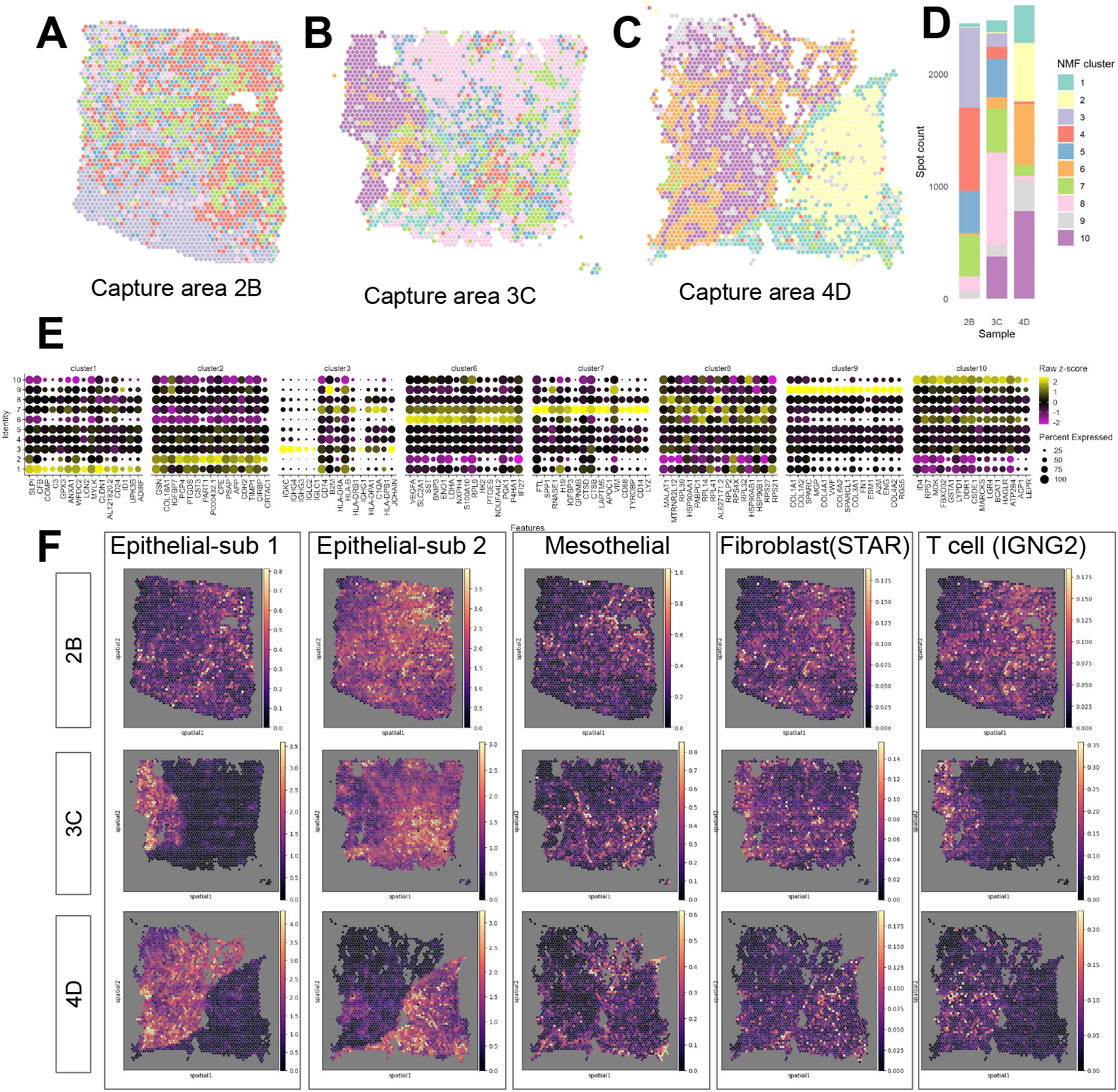
Characterizing intra-tumor heterogeneity using spatial transcriptomics. **(A-C)** Spot plot of predicted spatial domains (colors) using the R/Bioconductor package *ccfindR* [19] for *N* =3 tissue sections (capture areas 2B, 3C, 4D) from a single HGSC tumor. **(D)** Stacked bar plot summarizing the proportions of predicted spatial domains across the three tissue sections. **(E)** Dot plot displaying expression (each scaled by standard deviation and centered at 0) of di!erentially expressed genes (DEGs) for the predicted spatial domains using Seurat [20] with default parameters. NMF clusters 4 and 5 had fewer than 5 significant DEGs and hence were omitted. **(F)** Visualization of *Cell2location* [21] prediction of the 5^*th*^ percentile of selected cell type abundance for 3 capture areas (rows).

Next, we identified differentially expressed genes (DEGs) across the predicted spatial domains (**Figures 1E**). For example, in NMF cluster 1, we found DEGs for established ovarian cancer biomarkers, including *WFDC2* and *SLPI* [22, 23]. In contrast, in NMF cluster 3, which primarily comprised spots from sample 2B with smaller representation from 3C and 4D, showed upregulation in genes associated with B-cell or plasmablast activity, such as *IGKC, IGHG3, IGHG4*, and *JCHAIN*. In NMF cluster 6, *VEGFA*, known to be associated with endothelial activity and vasculature development, and *S100A10*, a hallmark gene for cancer migration [24], along with *LDHA* and *ENO1*, were upregulated, consistent with the typical cancer metabolism characteristic [25, 26]. In NMF cluster 7, we found *SPP1*, which is known to be expressed by cancer cells and tumor-associated macrophages [27].

Overall, the capture areas 3C and 4D were more similar in terms of both shared spatial domains and DEGs, compared to 2B (**Figure 1**). Specifically, the spatial organization of domains in both 3C and 4D both had a large contiguous area mostly occupied by NMF cluster 6 and 10 spots. The Moran’s I value of the top DEGs from NMF clusters are also more consistent comparing samples 3C and 4D (**Figure S2**). We found that capture area 2B lacked spatial domains 8 and 10 and was rather characterized by a spatial domain (cluster 3), which were enriched for DEGs associated with immune activity and were less abundant in capture areas 3C and 4D (**Figures 1A**,**B**).

To further validate the similarity and differences between tissue slices suggested by NMF clustering, the Python package *Cell2location* [21] was used to estimate cell type abundance using an annotated scRNA-seq dataset that is mapped to the Visium capture areas (**Figures S3**,**S4**,**S5**). Cell2Location results correspond well to the predicted spatial domains obtained via NMF clustering, with two distinct scRNA-seq epithelial cell populations (epithelial-subcluster1 and epithelial-subcluster2) roughly occupying two distinct spatial domains in samples 3C and 4D, as indicated by high *Cell2location* 5^*th*^ percentile cell abundance estimates of epithelial-subcluster1 in NMF clusters 6 and 10, and higher cell abundance estimates of epithelial-subcluster2 in most other NMF clusters (**Figures 1F**,**S3**). Notably, the separation of epithelial-subcluster1 and epithelial-subcluster2 were absent in sample 2B (as well as spatial domain 10) (**Figure 1F**). Such intra-tumor heterogeneity observed may suggest the necessity for generating spatially resolved sequencing data on multiple capture areas from the same tumor for a more comprehensive representation of its microenvironment, consistent with previous findings [28].

### 2.3 Predicting HGSC tumor subtypes using single-cell and spatial transcriptomics

In this section, we aimed to assess the intra- and inter-tumor spatial heterogeneity with respect to the predicted HGSC subtypes. Towards this goal, we expanded beyond the initial *N* =3 Utah HGSC Visium samples (all from the same tumor) to integrate these samples with a public dataset from Denisenko et al. [18] containing *N* =8 HGSC tumors (one capture area per tumor, sample names SP1-SP8) for a total of *N* =11 Visium capture areas across *k*=4 known HGSC subtypes (based on TCGA subtypes): differentiated (DIF), immunoreactive (IMR), proliferative (PRO), and mesenchymal (MES) (**Figures 2A-B**).

**Figure 2:**
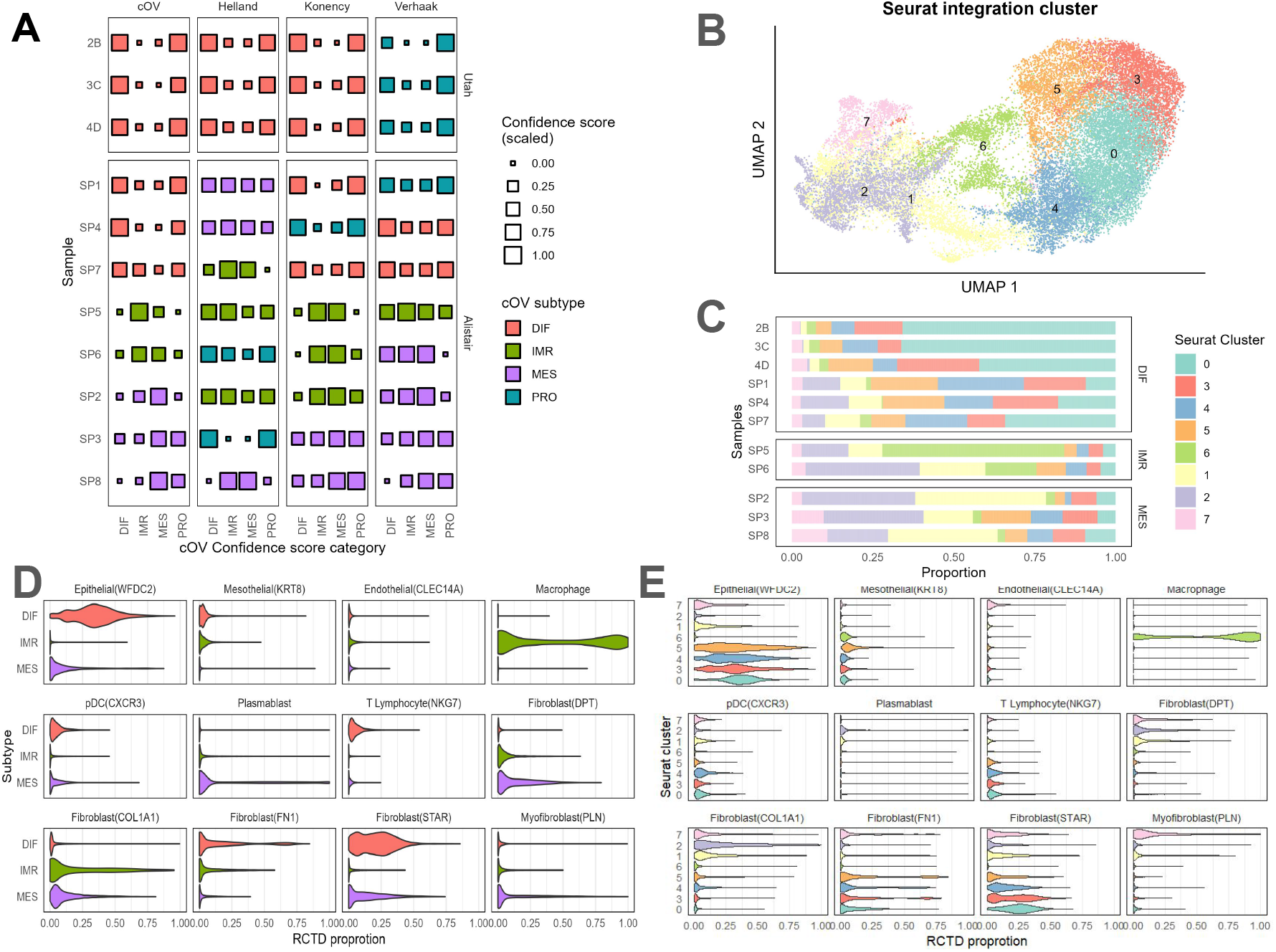
Characterizing inter-tumor heterogeneity using spatial transcriptomics. **(A)** Comparison of subtype assignment from 4 subtype prediction algorithms implemented by the *consensusOV* (cOV) R/Bioconductor package using pseudobulk RNA-seq profiles of *N* =3 Utah HCI Visium samples and *N* =8 Visium samples from [18]. **(B)** UMAP visualization of an integrated HGSC dataset (from *N* =11 HGSC capture areas). Colors represent the Seurat cluster of the integrated dataset. **(C)** Bar plot showing the spot-level predicted spatial domains for all *N* =11 Visium samples, with samples grouped by their predicted cOV HGSC subtype. **(D)** Violin plot showing the distribution of estimated cell types after applying spot-level deconvolution using CARD [29] for select cell types in the integrated dataset, aggregated over di!erentiated (DIF), mesenchymal (MES), and immunoreactive (IMR) pseudobulk cOV HGSC subtypes. **(E)** Violin plot showing distribution of estimated cell type proportions using CARD across the 8 predicted spatial domains.

We investigated whether we could predict the HGSC subtype using the aggregated molecular information across all spots in a given Visium capture area to a “pseudo-bulk” RNA-seq profile. We were interested in this task to assess if, for example, the spots from the *N* =3 capture areas (all from the same tumor) would be predicted to be the same HGSC subtype. For this, we used the *consensusOV* [9] R/Bioconductorv package to classify the HGSC tumors into *k*=4 subtypes. Within this package, we considered four different classifiers: (i) the consensusOV method (cOV), (ii) the Konecny method, (iii) the Verhaak method, and (iv) the Helland method [7–9, 30].

We found that the subtype predictions obtained using the cOV and Konecny methods demonstrated strong agreement with their final subtype assignment consistent in 8 out of the 11 capture areas. In contrast, the Verhaak method exhibited discrepancies with cOV and Konency method in six or more samples, while the Helland method produced different trends in confidence scores for samples SP7, SP6 and SP2 compared to cOV and Konency on top of differing subtype assignment (**Figure 2A**). Reassuringly, the 3 Utah Visium samples (2B, 3C and 4D) were consistently classified by all four prediction algorithms, with cOV, Konecny and Helland predicting DIF and Verhaak predicting PRO. Notably, the confidence scores assigned by all four methods assigned to the DIF and PRO subtype were both high and comparable for the three Utah samples, which according to the *consensusOV* manuscript, suggests ambiguity in classification potentially due to heterogeneous/admixed tumor tissue [9]. The Denisenko et al. [18] samples SP1, 4 and 7 were classified as the DIF subtype by the cOV method with high DIF confidence score. Interestingly, SP1 and 4 were also assigned high PRO confidence scores by the cOV method and directly classified as PRO by the Konency method [8]. Samples SP5 and SP6 were classified as the IMR subtype by the cOV and Konency methods. Finally, samples SP3 and SP8 were classified as the MES subtype consistently by the cOV and Konency methods, while SP2 was more ambiguously classified, predicted as MES by cOV and Verhaak and IMR by Helland and Konency (**Figure 2A, Supplementary Table S2**).

Next, using the same pseudobulked samples (with predicted HGSC subtypes), we assessed whether we would see similar predicted spatial domains at the spot level within a predicted HGSC subtype across the capture areas. To investigate this, we used both the predicted HGSC subtype labels and the spot-level integrated spatial transcriptomics datasets and visualized the intra- and inter-tumor spatial variation (**Figure 2C**). We found there were striking differences in the distribution of predicted spatial domains both within the same and across different predicted cOV HGSC subtypes (**Figure 2C**).

Then, we performed spatially-aware spot-level deconvolution for each of the *N* =11 capture areas using the Conditional Auto-Regressive-based Deconvolution (*CARD*) method [29] to estimate the proportion of cell types per spot. After applying CARD, we combined the samples based on the predicted HGSC subtypes using the cOV method to visualize the results. We found the *CARD* results demonstrated strong subtype-specific patterns similar to what is reported in a previously published study: specifically, DIF subtype spots were characterized by high proportions of epithelial cells, near-zero fibroblast, myofibroblast, and macrophage proportions, and low plasmablast proportions (**Figure 2D**) [5]. MES subtype spots exhibited mid to low proportions of epithelial cells, fibroblasts, and myofibroblasts, but remarkably high plasmablast proportions. The IMR subtype spots, on the other hand, were characterized by near-zero proportions of epithelial cells, fibroblasts, and myofibroblasts, mid to low plasmablast, and high macrophage proportions.

Finally, we visualized the estimated proportion of cell types from *CARD* stratified across predicted spatial domains obtained using Seurat (**Figure 2E**). Our hypothesis was that we would find consistent sets of cell types within a spatial domain. We found that the *CARD* results corroborated the subtype-domain association, with domains 0, 3, 4, and 5 (most abundant in the DIF samples) showing high proportions of malignant/epithelial cells and minimal fibroblast and macrophages. Conversely, clusters 1, 2 and 7 (most abundant in the MES samples) were marked by high fibroblast proportions and negligible epithelial cell presence. Interestingly, cluster 6, most abundant in IMR samples SP5 and SP6, exhibited high macrophage proportions, low epithelial, and low fibroblast proportions, and lacked plasmablast.

### 2.4 Functional characterization of predicted spatial domains in HGSC tumors

Next, we investigated biological functions underlying each of the predicted spatial domains within the HGSC TME. Specifically, we aimed to dissect the roles of spots enriched in epithelial, fibroblast, and immune cells by performing DGE analysis and, subsequently, gene set enrichment analysis on an integrated spot-level spatial transcriptomics dataset. To minimize the impact of batch effects due to the different experimental protocols used to generate the Utah Visium samples and the Denisenko et al. [18] Visium samples, in this section we did not use the Utah samples. Furthermore, we performed another integration step across only the Denisenko et al. [18] samples used in subsequent analyses.

Using unsupervised clustering, we obtained *k*=12 spatial domains after integrating the Denisenko et al. [18] samples (**Figures 3A**,**B**). We found that 5 spatial domains (0, 2, 7, 9, 10) had a high expression for a known marker for ovarian cancer *WFDC2* (**Figure 3C**). Therefore, we considered these domains to represent the epithelial-dominant parts of the tumor [31]. Five other spatial domains (3, 5, 6, 8, 11) were marked by upregulation of gene sets involved in extracellular matrix organization, extracellular structure organization, and collagen fibril organization. We annotated these domains as the stromal portion of the tumor (**Figures 3C**,**S6**). Domain 4 displayed DE gene sets associated with cytokine-mediated signaling pathways, antigen processing and presentation, innate immune responses, and several macrophage markers (*LYZ, CD68*) and was potentially representative of a mixture of infiltrating immune cells and tumor associated macrophages. Domain 1 had fewer DE genes and seemed to express moderate levels of both epithelial and stroma marker genes (**Figures 3C**,**S6**). Notably, spots in domain 1 were primarily localized between epithelial-dominant and fibroblast-dominant domains, further suggesting that it may represent a transitional area between the stroma and epithelial cells (**Figure S7**).

**Figure 3:**
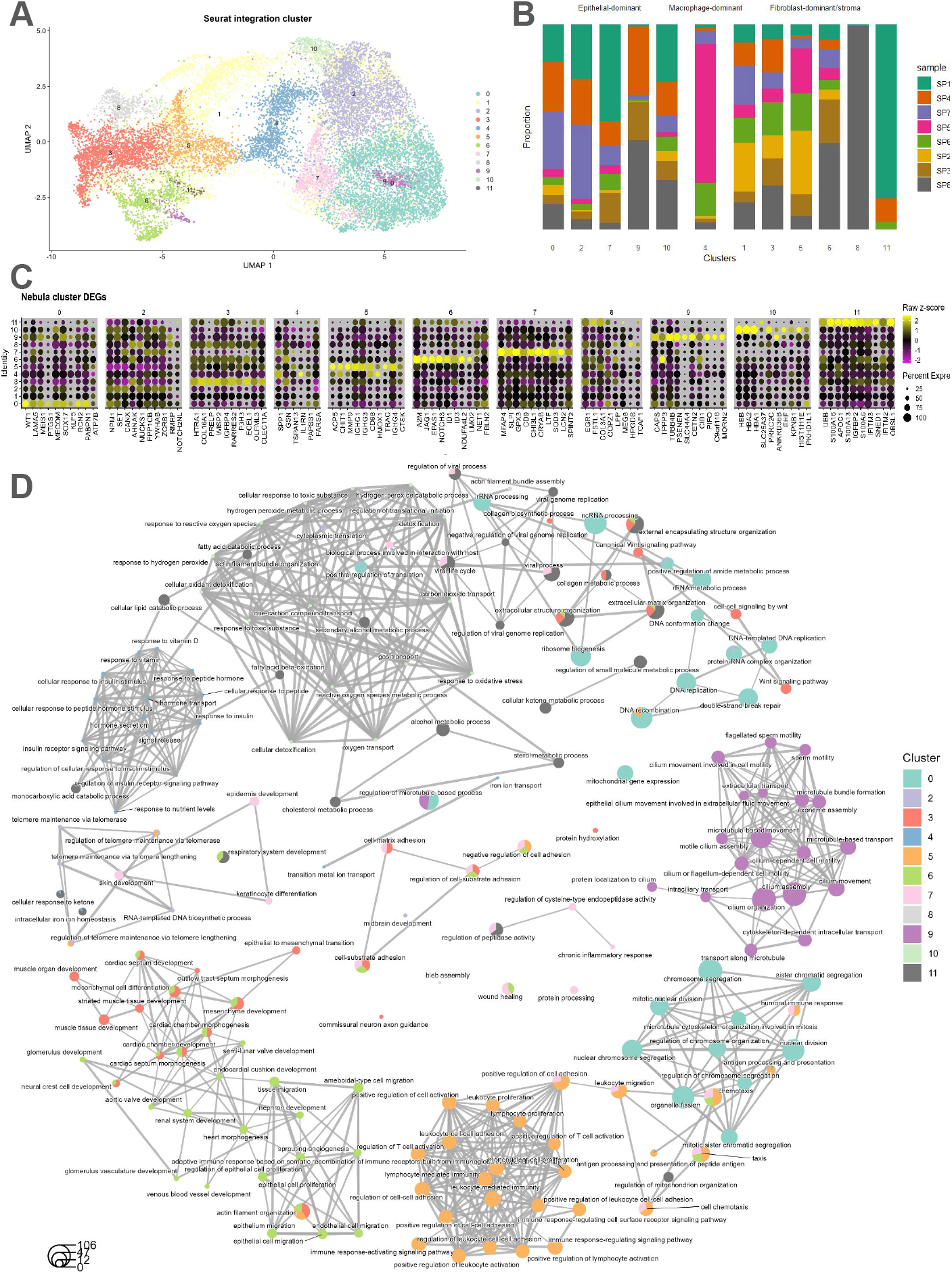
Functional characterization of predicted spatial domains in HGSC tumors. **(A)** UMAP representation of the *N* =8 Denisenko et al. [18] integrated Visium samples colored by *k*=12 predicted spatial domains. **(B)** Stacked bar plot representing the relative proportions of spots sourced from each sample stratified by each predicted spatial domain. **(C)** Dot plot of marker gene expression of cluster-specific DE genes. Spatial domain 1 had fewer than 5 significant DEGs and hence was omitted. **(D)** Gene concept network plot showing the GO (GeneOntology) results using *clusterProfiler* [32] based on spatial domain-specific, upregulated DEGs sets with the edges indicating the biological pathways.

#### 2.4.1 Functional characterization of epithelial cell-dominant cluster

To explore the biological functions specific to each epithelial cell-dominant cluster, we performed functional enrichment analysis using the filtered differentially expressed gene list (obtained after the removal of DEGs significant in more than one domain and corrected for cell composition).

The epithelial cell-dominant domain 0 exhibited upregulation of *LAMA5, SOX17, EFNB2*, which are known to be related to modulating cancer proliferation [33–35]. Domain 7 exhibited upregulation of *SLPI* and *CRYAB*, which are reported to promote cell proliferation and contribute to immune checkpoints [36, 37]. Domain 9 was characterized by upregulation of *CAPS*, a calcium binding protein essential for spindle formation, as well as tubulin subunits *TUBB4B, TUBA1A*. Furthermore, the DEGs from domain 9 also suggested functions pertaining to microtubule bundle formation, transport along microtubule and microtubule based movement (**Figure 3D**). Cluster 10 was marked by potential oxygen transport processes driven by upregulation of *HBB, HBA1*, and *HBA2* (**Figure 3C**).

#### 2.4.2 Functional characterization of fibroblast-dominant and stromal domain

Domain 3 was characterized by upregulation of collagen subunit *COL16A1* and *TNXB*, a protein regulating collagen assembly, as well as overexpression of *HTRA1* and *P3H3*, both reported as potential tumor suppressors in certain cancers [38, 39]. Domain 5 was characterized by B cell receptor signaling pathway and antigen receptor mediated signaling pathway involving upregulation of *IGKC, IGHG1, IGHG3*. Domain 6 was characterized by upregulation of *ID1* and *ID3*, DNA binding inhibitor proteins that are known to be required for vascularization in cancer tissue [40]. Interestingly, overexpression of *JAG1* and *NOTCH3*, a ligand receptor pair, was also observed in domain 6. Domain 11 was potentially involved with steroid, sterol, and cholesterol metabolic processes given upregulation of *SCARB1, UBB* and *APOC1* (**Figure 3C**).

#### 2.4.3 Biological theme comparison

Here, we analyzed enriched functional profiles of each of the gene clusters and aggregated the results into a single object. Comparing functional profiles can reveal functional consensus and differences among different experiments and helps in identifying differential functional modules in -omics datasets.

We found shared pathways by the stromal domains, such as extracellular matrix organization, collagen fibril organization, response to growth factor beta, as well as antigen presentation (**Figure 3D**). The epithelial-dominant domains were characterized by more distinct pathways: domain 0 and 9 both exhibited signatures of cellular respiration; domain 2 was characterized by functions such as regulation of protein localization to chromosome, telomere and positive regulation of DNA biosynthetic process; domain 7 was involved in growth factor response and antigen presentation; and domain 10 demonstrated oxygen transport and detoxification activities.

We generated biological theme comparison analyses of the genes whose expressions were positively associated with deconvolution-derived proportions. Functional annotations from genes associated with cell type abundance had a substantial overlap in basic biological processes like rRNA processing, ribosome biogenesis, and mRNA transcription (**Figure 3D**). Additionally, *KRT8* -expressing mesothelial cells were found to be enriched in functions such as protein polyubiquitination, while *COL1A1* expressing fibroblasts were functionally enriched in collagen biosynthetic and metabolic process. Interestingly, the functional annotations associated with immune cells except pDC are mostly descriptive of subcellular activities: endosome to melanosome transport and cellular pigment accumulation are the top biological themes for T lymphocytes (potentially reflective of cell proliferation in spots with higher T cell abundance); ER to cytosol transportation and cellular response to unfolded proteins are the top themes associated with spots of high plasmablasts abundance [41]; cross-membrane transport and ion homeostasis are the top theme for monocytes [42] (**Figure S9**).

### 2.5 Immunoreactive HGSC subtype areas have anti-correlation between immunoglobulin and major histocompatibility complex genes

To examine potential gene-gene interactions between clusters, we selected the top DEGs by log fold-change from each cluster, plus 2 isolated immune gene programs (histocompatibility and immunoglobulin DE genes) and calculated the correlation between their expression (**Figure 4**).

**Figure 4:**
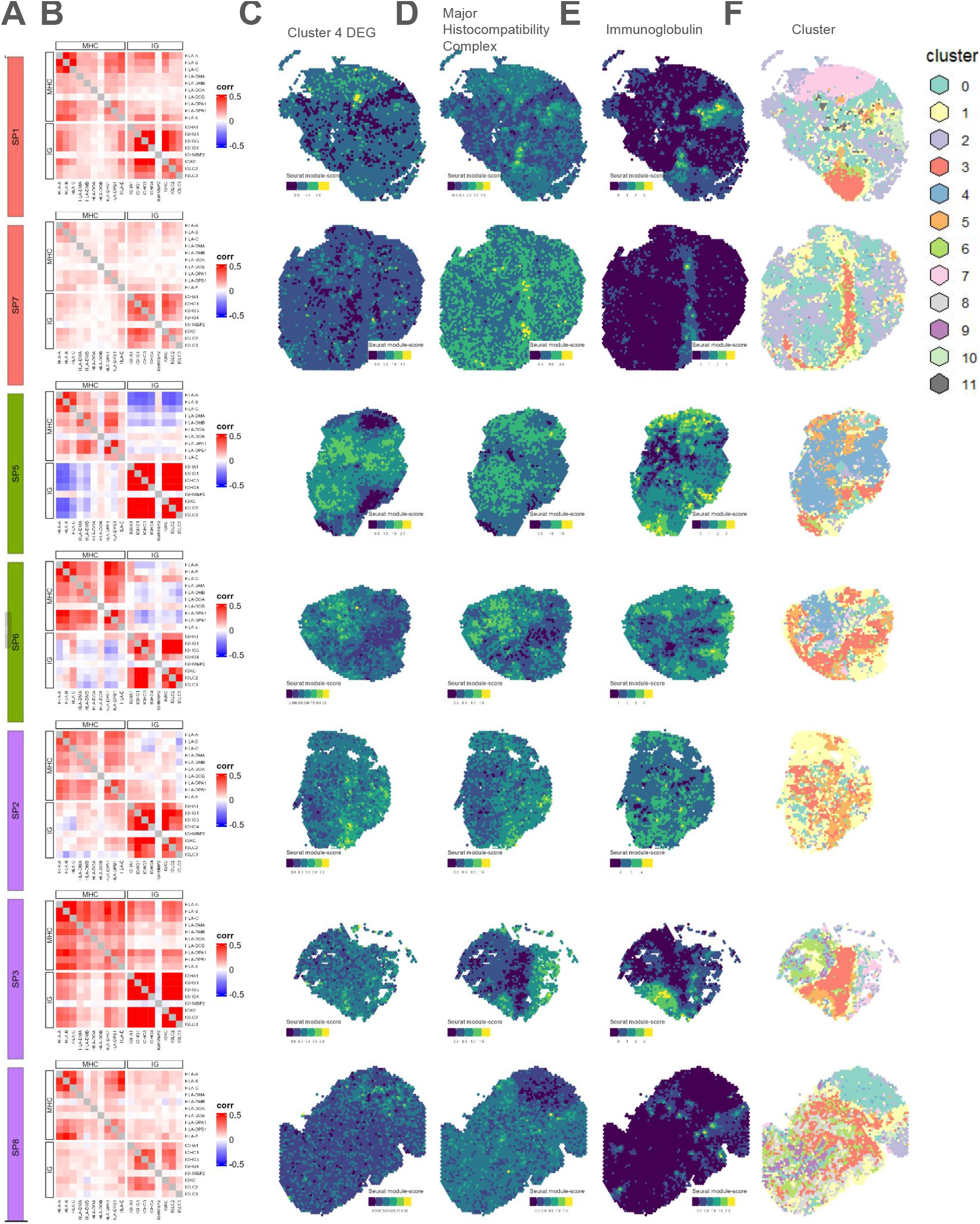
Immunoglobulin and major histocompatibility complex genes are anti-correlated in the immunoreactive subtype, but not in the di!erentiated or mesenchymal subtypes. Data visualization of predicted spatial domains using Seurat in select Visium samples from Denisenko et al. [18]. We do not include the spatial domain visualization for sample SP4 (dif) here, as it displays much less variability in spatial domains in comparison to other DIF samples. Please see **Figure S8** for SP4 spatial domains. **(A)** Sample id and colorcoded molecular subtype (di!erentiated = orange, immunoreactive = green, mesenchymal = purple). **(B)** Spearman correlation values between domain 4 DEGs, major histocompatibility complex (MHC) genes and immunoglobulin (IG) genes. There was a negative correlation between MHC and IG exclusively for the immunoreactive samples. Column corresponding to the Seurat module score of DEGs from identified from domain 4. **(D)** Column corresponding to the *Seurat* module score (average gene program expression) of major histocompatibility complex coding genes. **(E)** Column corresponding to the Seurat module score of immunoglobulin coding genes. Note: spots tend to co-express high MHC and IG in the DIF and MES samples (top and bottom 2 rows) while spots expressing high MHC tend to anti-colocalize with spots expressing high IG in IMR samples (rows color coded green). **(F)** Integrated clustering scheme in space for reference.

We found that the correlation patterns among DEG sets varied across the *N* =8 Denisenko et al. [18] Visium samples. In particular, the major histocompatibility complex (MHC) and immunoglobulin coding genes demonstrated the most apparent subtype-specific interaction (**Figures 4A-B**). In the samples with a DIF subtype or MES subtype, we found zero to weak positive Spearman correlations between these two gene sets. Interestingly, spatial variability in these gene sets that is observed in other DIF subtype samples sample is not apparent in SP4 (DIF) **Figure S1**. Expression of these MHC and immunoglobulin coding genes, if both were expressed, also co-localized in space (**Figures 4C-F**). In samples with the IMR subtype, we found a negative Spearman correlation between type-II MHC and immunoglobulin coding genes (**Figure 4B**), and expression of the two immune gene sets seems to anti-co-localize in space (**Figures 4C-F**). The separation of immunoglobulin expression from histocompatibility antigen expression, which seemed to exclusively occur in the IMR subtype samples, may indicate plasmablast differentiation and movement away from the site of antigen presentation.

### 2.6 Investigating cell-cell communication in HGSC tumors across spatial domains

Lastly, we utilized the COMMunication analysis by Optimal Transport (COMMOT) [43] package to predict cell-cell communication through secreted signaling and to detect differences in tumor tissues at the molecular subtype level and beyond (**Figures S10A-B**). A total of 172 ligand-receptor pairs from the CellchatDB database [44] were found to be expressed in at least five spots across all [18] samples. The top cell-cell communication pathways detected across all samples include MK (*MDK* -*SDC1, MDK* -*SDC2, MDK* -*SDC4, MDK* -*LRP1, MDK* -*NCL*), MIF (*MIF* -*CD74* -*CXCR4* and *MIF* -*CD74* -*CD44* (**Figure S10B**). Pathways CXCL (*CXCL12* -*CXCR4*), COMPLEMENT (*C3* -*C3A41, C3* -*ITGAM* -*ITGB2* and *C3* -*ITGAX* -*ITGB2*) interactions are highly active in samples of the immunoreactive subtypes (**Figure S10B**). This aligns with previous findings on CAFs and ovarian cancer cell interactions reported by [45]. The COMMOT-derived aggregate estimates for outgoing signals across these pairs generally exhibited a bell-shaped distribution. The mean aggregated signal levels were remarkably consistent within the same molecular subtype, with the IMR subtype exhibiting the highest mean signal levels, followed by the MES subtype. In contrast, differentiated spots showed significantly lower mean signals and a narrower distribution (**Figure S10A**).

We then examined the correlation between ligand-receptor pairs, and found subtype-specific correlation patterns. Most notably, the high within-pathway correlation of the MK pathway in sample SP5 suggests the lack of differential colocalization of *MDK* mediated signaling in space. By contrast, in samples SP1 and SP7, where *MDK* -*LRP1, MDK* -*NCL* and *MDK* -*SDC4* signaling colocalize to different clusters, there is little intra-MK correlation. The within-pathway correlation of the MIF pathway is relatively high for immunoreactive samples SP5 and SP6 as well as mesenchymal sample SP2 and much less in the differentiated samples SP1, SP4 and SP7 (**Figure S11**).

#### 2.6.1 Differences in ligand-receptor activity between molecular subtypes

Further analysis of cluster-specific signal distribution revealed that fibroblast-dominant clusters 3, 5, and 6 had the highest levels of outgoing signals. In non-DIF samples, the epithelial-dominant cluster 0 also showed high signaling activity, notably more so than cluster 1, despite their frequent spatial proximity (**Figures S10B-C, S7**). In DIF samples (SP1, SP4, and SP7), clusters 1 and 0 displayed similar signal levels, much lower than those in clusters 3, 5, and 6, suggesting reduced secreted signaling interactions in epithelial-dominant clusters of DIF samples.

#### 2.6.2 Ligand-receptor signaling occurs in different spatial domains across HGSC samples within the DIF subtype

Midkine-Syndecan-1 (*MDK* -*SDC1*) signaling is virtually globally active in SP1 while only weakly present in specific clusters (5, 7) of SP4 and SP7. The Midkine-Syndecan 4 signaling is active in SP4 and SP7 in a global manner, while only prominent in cluster 7 of SP1 (**Figures 5A-B**). This observed differential regulation of Syndecan-1 and 4 across the differentiated subtype samples may indicate distinct functions underlying these ligand-receptor interactions, and is potentially consistent with the results from a previous study on breast carcinoma, which reported significant association between Synedcan-1-positive staining and Synedcan-4-negative staining with tumor grade suggesting they are independent indicators in breast cancer [46].

**Figure 5:**
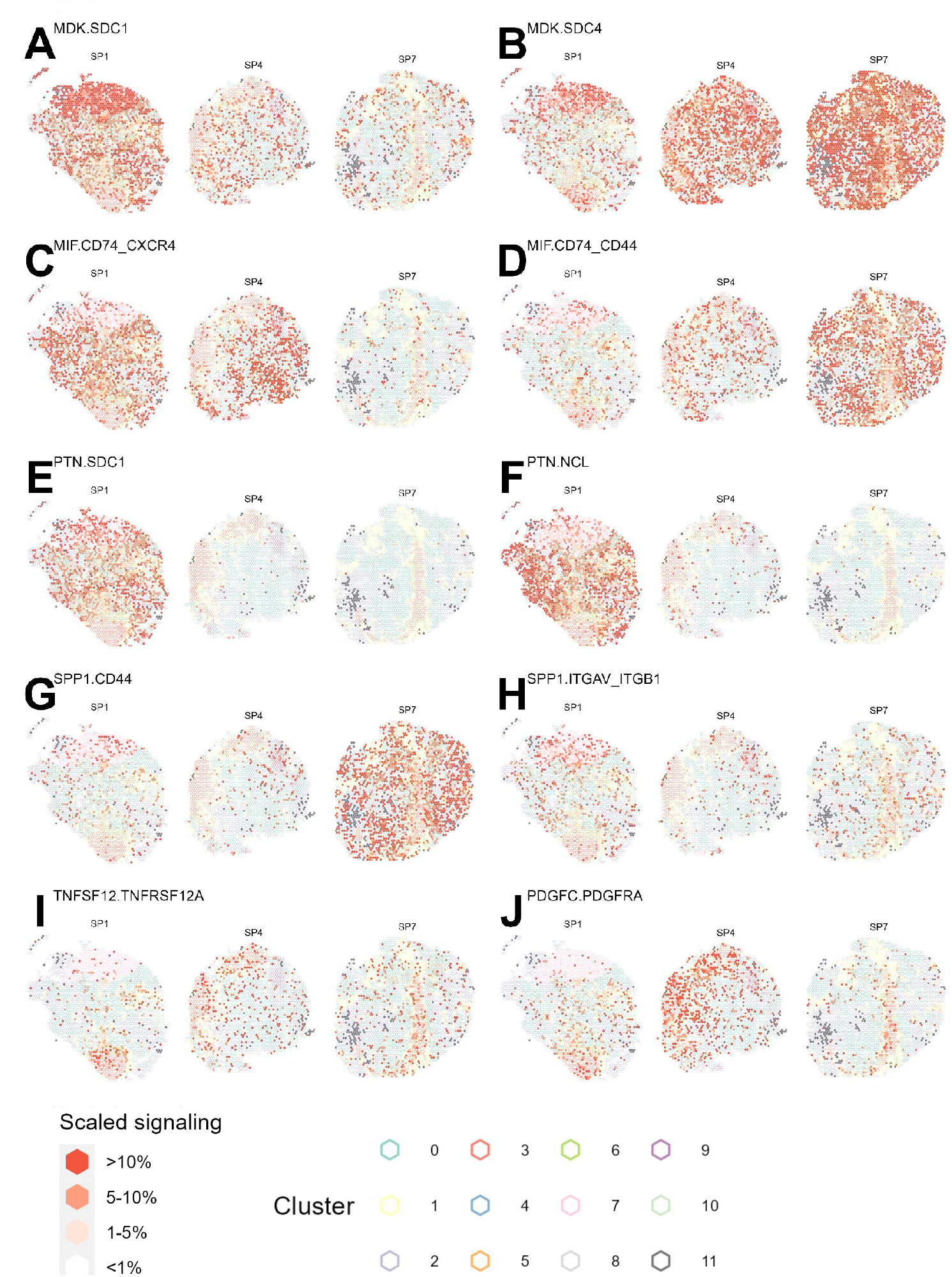
Ligand-receptor signaling occurring in different domains across samples of the DIF subtype. Each panel **(A-I)** displays the outgoing signal strength predicted by COMMOT for a different ligand-receptor pair in 3 DIF samples SP1, SP4 and SP7, respectively. The fill of the spots represents binned signaling across all spots; the outer boundary are color-coded by the spatial domain.

*MIF* -*CD74* -*CXCR4*, the non-cognate ligand receptor interaction that is reported to have chemokine-like functions and result in recruitment of immune cells, has prominent signaling present in the epithelial cell rich clusters of SP1 and SP4 (clusters 0, 9, 10, 11) while much lower in SP7. *MIF* -*CD74* -*CD44*, the canonical interaction that triggers the standard ERK MAP kinase pathway activation, is substantially upregulated in SP7 while only present in clusters 5, 6, 7 in SP1 and SP4 [47] (**Figure 5C-D**). A recent study featuring reported cell-cell communication between macrophages and T cells in ovarian cancer via the *MIF* -*CD74* -*CXCR4* pathway altering CD8-T cell function and impacting prognosis [48]. This difference in MIF signaling observed within the DIF subtype is potentially reflective of the differing immune cell state between SP7 and the other DIF samples, especially under the observation that SP7 had a substantially higher proportion of spots of the immune cell-dominant cluster 4 (**Figure 3B**).

*PTN* is known as a mitogenic cytokine whose signaling is associated with neural development as well as various cancer related biological activities [49]. Interestingly, PTN signaling appears exclusively in the epithelial cell-dominant clusters (0, 7, 10) of sample SP1 and no other sample (**Figure 5E-F**).

*SPP1* -*CD44*, a pathway known to carry out immunosuppressive functions [50], is globally active in SP7 and absent from the two other DIF samples. *SPP1* -*ITGAV* -*ITGB1* interaction, which is reported to promote tumor progression in ovarian cancer [51], is present in all 3 DIF samples in moderate levels and yet appears to colocalize with different clusters. In SP1, *SPP1* -*ITGAV* -*ITGB1* signaling happens primarily along the boundary lines of cluster 7 as well as along the division of clusters 0 and 1; In SP4, *SPP1* -*ITGAV* -*ITGB1* colocalizes almost exclusively with cluster 9; In SP7 *SPP1* -*ITGAV* -*ITGB1* is most prominent along fibroblast-dominant clusters 3, 5, 6 and cluster 7 (**Figure 5G-H, S12, Supplementary Table S3**).

The TWEAK-Fn14 (*TNFSF12* -*TNFRSF12A*) interaction, known to activate cellular processes such as proliferation, invasion, and angiogenesis in cancer [52], is present in all 3 DIF samples in moderate levels and colocalizes primarily with fibroblast rich cluster 3. In SP4 and SP7, this interaction is additionally observed in the epithelial cell-dominant clusters 0 and 7 (**Figure 5I**).

Such differences in cell-cell communication pathways between DIF samples are indicative of the heterogeneity of tumors within the sample’s molecular subtype. Most interestingly, differences in signaling patterns include both global differences in signaling strength and the specific co-localization of signaling with spatial domains, further highlighting the presence of unique tumor architectures.

## 3 Discussion

Utilizing spatial transcriptomic data from several sections from the same tumor, we explored intra-tumor heterogeneity. We also expanded this analysis to intra- and inter-tumor heterogeneity by integrating publicly available HGSC tumors.

Our findings on the variability of the Visium dataset taken from the same tumor sample is potentially suggestive of the limited ability for a single Visium tissue slice to comprehensively characterize the complex TME and calls for the need to generate multiple Visium slices. However, a technical limitation underlying our Visium samples is that we sequenced fresh frozen tissue, which may result in undesired tissue stretching and is more vulnerable to poor quality tissue from RNA degradation compared to formalin-fixed paraffinembedded (FFPE). Replication of these findings using FFPE tissue will be needed to better validate our observed patterns of intra-tumor heterogeneity.

The application of spatially-aware deconvolution, using the *CARD* package, with a molecular subtype classifier, *consensusOV*, on the pseudobulk of Visium samples allowed us to identify a connection between tumor composition and molecular subtype. Our analysis of spatially resolved transcriptomic data confirm the significance of molecular subtypes of HGSC in spite of the existence of considerable inter-tumor variability and complexity of solid tumor tissue.

Subsequent functional enrichment analysis on the integrated Visium samples revealed spatial domains with distinct biological functions shared across tumor tissues. Further using spot-wise Spearman correlation between expression of DEGs, we detected an immunoreactive subtype-specific negative interaction between MHC genes and immunoglobulin genes, which appears to be a novel finding and may indicate potential interaction of plasma cells and antigen-presenting cells. One possible explanation of this phenomenon is immunoreactive subtype-specific movement of B cells away from the site of antigen presentation, which may indicate stronger or later-stage immune infiltration. However, we only analyzed two immunoreactive samples in this manuscript, which limits our ability to arrive at concrete conclusions. Further research incorporating more HGSC samples with reliable molecular subtype classification will be needed to validate whether these interactions are prevalent in and specific to immunoreactive samples. Moreover, spatially resolved transcriptomic technology at single-cell resolution, such as Vizgen MERFISH and 10x Genomics Xenium and can be leveraged to interrogate the spatial organization of plasma cell and antigen-presenting cells and confirm whether an anti-co-localization occurs in immunoreactive samples.

Lastly, we interrogated the cell-cell communication patterns of all samples across all DIF subtype samples. Our application of COMMOT, a spatially-aware method based on optimal transport for cellcell communication analysis, has unveiled considerable heterogeneity in signaling activity across tumors of different molecular subtypes. The differential colocalization of signaling activities to distinct spatial domains within each tumor could be a driving factor for recruitment of cells and thus shaping the TME. We identified potential cell-cell communication features we could extract from spatially resolved data, such as strength of correlation between immune gene sets and presence/absence of signaling pathways, that may be valuable covariates for consideration when assessing prognosis. Potential future directions include further analysis of spatially resolved data searching for marker gene expression strong associated with such spatial features to enable survival analysis population level using bulk-RNA-seq as input.

Also quite interestingly, despite analyzing the same ovarian high grade serous carcinoma dataset published by [18], our cell-cell communication results are hardly comparable due to the drastically different methodology being used (the Denisenko et al. [18] study focused on the correlation between expression of select ligand and Giotto cell type abundance scores in a patient-specific manner, while the COMMOT package takes into account the spatial information and potential heterodimeric interactions).

A major limitation of the study, on top of the small amount of samples used in the analysis, was the lack of incorporation of patient-level metadata on therapy and survival information. Analysis of more spatially resolved transcriptomics data with inclusion of patient level metadata should be done to verify our current findings and explore additional features in spatial data associated with treatment and survival. Additionally, the cell-cell communication analysis is subject to the limitation of having a pre-defined communication distance threshold (`dis_thr`) parameter. We selected a relatively small value of 250 for all samples, which only allowed for the detection of short distance secreted signaling.

In conclusion, our study has introduced novel approaches for analyzing single-cell and spatially resolved transcriptomics data through integrating molecular subtyping information with correlation of differential expression sets and cell-cell communication. Our results suggest potentially useful features that can be extracted from spatially resolved transcriptomics data and be applied to population level analysis of HGSC.

## 4 Methods

### 4.1 HGSC tumor tissue sample and tissue processing

#### Utah Visium data generation

One fresh-frozen tumor sample (JD24379) was shipped from the Huntsman Cancer Institute at the University of Utah to the Cedars-Sinai Applied Genomics, Computation & Translational Core on dry ice. The tumor was embedded in Optimal Cutting Temperature (OCT) compound per the demonstrated protocol for tissue preparation (Demonstrated Protocol, CG000240). For RNA quality assessment, ten sections of 10*µ*m thickness were collected in a pre-cooled microcentrifuge tube, transported on dry ice, and stored at -80°C per the demonstrated protocol. RNA extraction was performed using the Zymo Quick RNA micro kit, and RNA integrity was evaluated on the 2100 Bioanalyzer. Methanol fixation, H&E staining, and imaging were performed per the demonstrated protocol (Demonstrated Protocol, CG000160). Tissue optimization was performed following 10X Genomics Visium Spatial for Fresh Frozen Tissue Optimization protocol (User Guide, CG000238). Spatial gene expression libraries were generated following 10X Genomics Visium Spatial Gene Expression for Fresh Frozen protocol (User Guide, CG000239). Libraries were sequenced on an Illumina NovaSeq 6000 instrument. Length of read 1 was 28 bp, i7 Index and i5 Index were 10 bp, and read 2 was 90 bp.

#### Utah scRNA-seq data generation

Samples were collected from seven patients with advanced stage (III or IV) ovarian high grade serous carcinoma (HGSC) by the Huntsman Cancer Institute at the University of Utah. Five patients underwent primary debulking surgery prior to adjuvant chemotherapy, and two patients (corresponding to samples 19833X1 and 19833X2) received 3 cycles of neoadjuvant chemotherapy prior to surgery followed by adjuvant chemotherapy.

In preparation for single cell RNA-seq, fresh macrodissected tumor samples (19459X1, 19595X1, 19833X1, 19833X2) were cut into 2 mm chunks and dissociated using the Human Tumor Dissociation kit from Miltenyi Biotec (130-095-929), according to the manufacturer’s instructions, in C tubes (130-093-237) with RPMI (ThermoFisher Scientific 11875135) + 10% fetal bovine serum (FBS, ThermoFisher Scientific 26140079). Dissociation was done on gentleMACS Octo dissociator from Miltenyi Biotec at 37°C for 1 hour. The cellular suspension was filtered using a 70 micron cell strainer (Miltenyi Biotec 130-098-462). Filtered cells were washed twice using a PBS + 0.04% FBS solution. Washed cells were diluted to obtain a concentration of 600,000-1,400,000 live cells/mL, followed by cryopreservation in liquid nitrogen.

Thawed dissociated cell suspensions were partitioned into an emulsion of nanoliter-sized droplets using a 10X Genomics Chromium Controller and Chromium Next GEM Chip G, and RNA sequencing libraries were constructed using the 10X Genomics Chromium Next GEM Single Cell 3’ Reagent Kit v3.1. Purified cDNA libraries were qualified on an Agilent Technologies 2200 TapeStation using a D1000 ScreenTape assay (cat# 5067-5582). The molarity of adapter-modified molecules was determined by quantitative PCR using the Kapa Biostystems Kapa Library Quantification Kits, KK4824 (cat# 0796014001). Individual libraries were normalized to 5 nM and equal volumes were pooled in preparation for Illumina sequence analysis. Sequencing libraries were sequenced on an Illumina NovaSeq 6000 instrument using the NovaSeq 6000 S4 v1.5 reagent kit (cat# 20028312; paired-end, 150x150bp).

Experimental details for samples 16030X2, 16030X3, and 16030X4 were previously reported [53], and data is available through dbGaP (accession phs002262.v2.p1). Briefly, tumor chunks cryopreserved in liquid nitrogen were thawed and dissociated into single cells using the Miltenyi Human Tumor Dissociation Kit and the GentleMACS dissociator. Library preparation was performed using the 10x Genomics 3’ Gene Expression Library Prep v3. Libraries were sequenced on an Illumina NovaSeq 6000 instrument.

### 4.2 Data processing and quality control

For all analysis steps described, we used Python (version 3.7), R (version 4.3.1), Seurat (version 5.0.1), and Bioconductor (version 3.18) for analysis of genomics data. SpaceRanger output was read into Python to create an AnnData object and read into Seurat and Bioconductor to create SeuratObjects and SpatialExperiment objects [20, 54].

#### Utah Visium raw data processing and quality control

For the tumor sample (JD24379), FASTQ and image data were pre-processed with the 10x SpaceRanger pipeline version 3.0.1. Reads were aligned to reference genome GRCh38-2020-A. Seurat S3 objects of version 4.1.3 were constructed for the four Visium capture area of the tumor sample, with subsequent analysis performed with Seurat version 4.3.1.

One out of four Utah HCI Visium samples had considerably fewer read counts mapped to each spot (3000 → 5000) compared to the others and was hence excluded from downstream analysis. For the other Utah Visium samples, we did not remove any spots based on standard quality control metrics including percent of reads mapping to mitochondrial genes. After applying these QC steps (removal of capture Area 1A), the number of gene by the number of spots was 18162 genes x 7537 spots.

Raw counts were normalized using Seurat `SCTransformation`, regressing out percent of mitochondrial genes, percent of ribosomal genes, and total UMI counts.

#### Utah scRNA-seq raw data processing and quality control

Following sequencing, we mapped reads to GRCh38-2020-A using cell ranger (version 7.0.0). After alignment, we perform quality control and dropped only cells expressing fewer than 200 genes or more than 25% mitochondrial genes. Genes expressed in fewer than 3 cells were also excluded. The dimensions of our dataset pre/post cell-level quality control are detailed in **Supplementary Table S1**.

#### Denisenko Visium data processing and quality control

Denisenko et al. [18] made available the standard outputs from 10x Genomics `Space Ranger and Loupe Browser` (filtered matrix, scalefactor and tissue positions files) for *N* =8 Visium datasets, which were used to construct Seurat S3 objects of version 4.1.3. However, subsequent analysis was performed with Seurat version 4.3.1.

For the Denisenko et al. [18] Visium samples, we did not remove any spots based on standard quality control metrics including percent of reads mapping to mitochondrial genes, total number of UMI counts per spot, or number of detected genes. This resulted in a total number of 15193 genes x 20123 spots.

Raw counts were normalized using Seurat `SCTransformation`, regressing out percent of mitochondrial genes, percent of ribosomal genes, and total UMI counts.

#### Denisenko scRNA-seq raw data processing and quality control

Denisenko et al. [18] made available the count matrices (in tab delimited file format) of the five scRNA-seq samples (Y2, Y3, Y5, MJ10, MJ11). For these scRNA-seq samples, cells expressing fewer than 200 genes or more than 25% mitochondrial genes were removed. Genes expressed in fewer than 3 cells were also excluded. The dimensions of our dataset pre/post are detailed in **Supplementary Table S4**.

### 4.3 Data analysis

#### Feature selection, dimensionality reduction, and batch effect correction for Visium samples

Features (genes) were selected using the *Seurat* `VariableFeatures` function using default parameters for all individual SRT samples, variable genes identified in at least 2 samples were selected as integration features. Following feature selection, principal component analysis (PCA) was performed with Seurat `RunPCA` with `npcs` = 30. To visualize the spots from the integrated dataset, we generated uniform manifold approximation (UMAP) representation of the principal components.

To correct for batch effects, we integrated the spots across the Visium samples using `SelectIntegrationFeatures` function in *Seurat* with 3000 features, `FindIntegrationAnchors` under reference = NULL and reduction set as `cca`, and `IntegrateData` using the 30 principal component dimensions for anchor weighting procedure while explicitly setting `features.to.integrate` as all genes.

Unsupervised clustering to identify predicted spatial domains for Utah Visium samples. To predict spatial domains, we used non-negative matrix factorization (NMF). Specifically, the variable Bayes NMF function vb factorize from *ccfindR* package was run on the merged raw count matrices of spatially variable genes from the *N* =3 Visium Utah samples, over ranks ranging from 8 to 16, with 5 replicates per rank [19]. The algorithm identify an optimal rank of 10 and assigned spots from all 3 samples into 10 predicted spatial domains.

#### Unsupervised clustering to identify clusters in the integrated Visium dataset

Following integration with *Seurat* `IntegrateData` as described previously, we ran the *Seurat*`FindNeighbors`using all 30 pricipal components and then the `FindClusters`() function at resolution 0.4 and default **Louvain** algorithm to obtain the clusters for (the same clustering methodology was applied to both Utah, Alistair/Denisenko et al. SRT integration and Denisenko et al. SRT integration).

#### Differential gene expression for Utah Visium sample across NMF predicted spatial domains

Preliminary differential expression analysis was performed on the integrated dataset using Seurat `FindMarkers`function under `log fold-change.threshold`parameter of 0.25 and `min.pct parameter`of 0.1, comparing spots from each NMF cluster (with more than 50 spots) with all other spots (in a one vs all manner).

*Nebula* was used for the formal differential expression analysis. The integrated`SeuratObject`of the 3 Utah samples were supplied to the *Nebula* `scToNeb`function, with the assay set as Spatial (raw counts), clusters as predictors and offset set as UMI number per spot (`nCount Spatial`), to generate the *Nebula* object. The design matrix supplied to the *Nebula* command included NMF cluster (one hot encoded, the first cluster with less than 5 differentially expressed genes from the preliminary differential expression analysis and are set as the reference cluster) and CARD deconvolution proportions for all cell types. Entries in the *Nebula* results that failed to converge (convergence indicator ↑ →20) are discarded. A gene was deemed differentially expressed in a cluster if that cluster was associated with the highest statistically significant log fold-change coefficient.

#### Differential gene expression for Denisenko Visium dataset and downstream analysis

Preliminary differential expression analysis for the integrated Denisenko Visium dataset was performed using *Seurat* `FindMarkers`function under log `fold-change.threshold`parameter of 0.25 and `min.pct parameter`of 0.1, comparing spots from each cluster with all other spots (one vs all).

Nebula was used for the second round differential expression analysis. The integrated Seurat object of the 8 Denisenko samples were supplied to the *Nebula* scToNeb function, with the assay set as Spatial (raw counts), clusters as predictors and offset set as the UMI number per spot (`nCount Spatial`), to generate the *Nebula* object. The design matrix supplied to the Nebula command included Seurat cluster (one hot encoded, the first cluster with less than 5 differentially expressed genes from the preliminary differential expression analysis and are set as the reference cluster) and `CARD`deconvolution proportions for all cell types. Entries in the *Nebula* results that have failed to converge (convergence indicator ↑ →20) were discarded. A gene was deemed differentially expressed in a cluster if that cluster was associated with the highest statistically significant log fold-change coefficient.

After obtaining the DEG sets corresponding to each cluster, we isolated two immune gene programs (histocompatibility and immunoglobulin DEGs) and calculated the Spearman correlation across sets of DEGs to quantify the degree to which upregulated cluster-specific gene sets interact. The results were visualized using heatmap.

#### snRNA-seq feature selection, dimensionality reduction, and unsupervised clustering

Integration of Utah and Denisenko single-cell datasets was performed with *scVI-tools* v1.0.4, *scanpy* v1.9.6, *anndata* 0.1.3. Pre-integration feature selection of top 3000 highly variable genes was performed on merged raw count matrices using *scanpy* with flavor parameter set as `Seurat v3`, and `batch key`parameter set as sample identifier [55–57]. Additionally, the marker genes from 32 stromal subclusters from Qian et al. [58] Pancancer blueprint were included as the features to integrate. Integration was achieved by training a scVI model with 2 layers, latent space dimension of 30, and negative binomial likelihood model and parameter `batch key`set as sample identifier.

Clustering of *scVI* [55] integrated single-cell dataset was performed with *scanpy* `pp.neighbors`and `tl.leiden`function with default parameters, with the *scVI* learned latent representation, resulting in 19 total clusters (**Figure S4A**).

#### Differential gene expression using scVI integrated scRNA-seq data

Differential expression analysis for the integrated single cell was performed with *scVI-tools* differential expression function, using leiden clusters as grouping variables (and by default conducts 1 cluster vs all testing). The DEGs were filtered by `lfc mean`*>* 2.0, bayes factor *>* 3, non zeros proportion1 *>* 0.5 and non zeros proportion2 *<* 0.25

Three clusters (clusters 0, 1 4 and 6) had less than 10 DEGs under such criteria. Four of the 19 leiden clusters (cluster 5, 6, 8, 9) were identified as tumor/epithelial cells due to their expression of canonical epithelial ovarian cancer markers including *WFDC2*, and *ELF3* (**Figure S5A-B**) [31, 59]. Clusters 12 and 16 were identified as mesothelial cells by marker gene expression of *CALB2, KRT8, KRT18* and *WT1* [11]. Cluster 7 was identified as endothelial cells by expression of canonical marker genes *CLDN5, VWF* and *CLEC14A* (**Figure S5A-B**) [11, 58].

Seven of the 19 leiden clusters were identified as immune cells: First, a T/NK cell cluster (2) expressing *NKG7* and *CD3D* was detected [58]. Another cluster (10) expressing *FKBP51* and *RHOH* appeared as an additional T lymphocyte cluster [60] [61]. Cluster 15 expressed *IGHG1* and *IGHG4* and was denoted B/Plasmablast. Cluster 3 showed expression of *CD68, MSR1* and *MRC1* and was hence identified as macrophage. Cluster 17 expressed considerably less macrophage marker *CD83* and showed high expression of *CD14* hence we denoted it as a separate monocyte population. Cluster 18 lacked *CD68* but expressed high levels of CD83 as well as pDC markers *LILRA4* and *CXCR3* and hence was identified as a pDC population (**Figure S5A-B**) [58].

Six of the 19 leiden clusters were identified as Fibroblast or Myofibroblasts: Cluster 13 showed high expression of *LUM, DCN*, and *DPT* and was denoted as Fibroblast (DPT) population. Clusters 11 and 16 showed expression of college subunits including *COL1A1, COL1A2* and *COL3A1*, with cluster 16 additionally expressing *FN1* and *EBF1*, and hence were classified as Fibroblast (COL1A1) and Fibroblast (FN1), respectively. Clusters 0 and 1 had moderate expression of *DCN* and expressed moderate to high amounts of *STAR* and *FOXL2* gene and were identified as two Fibroblast (STAR) populations. Lastly, cluster 4 expressed canonical myofibroblast marker genes *ACTA2* and *TAGLN*, hence was denoted as Myofibroblasts (**Figure S5A-B**) [58].

Upon examining the expression of marker gene sets published by [18], we were able to recover most celltypes (Tumor/Epithelial, Mesothelial, Endothelial, T cell, B cell, Macrophage, Fibroblast subpopulations 1, 2, 3, 5 and Myofibroblasts). The profile of Denisenko et al. [18] Fibroblast 4, marked by expression of *CCL2*, did not match any distinct leiden cluster in our integration. Nonetheless, the Myofibroblast population (leiden cluster 4) in our integration had moderate expression of *CCL2*, potentially indicating that some of the cells corresponding to the Denisenko et al. [18] Fibroblast 4 populations may be clustered with Myofibroblast by scVI integration (**Figure S5C**).

### 4.4 Functional enrichment analyses

Functional enrichment analysis comparing functional profile of clusters was performed with `compareCluster`and `enrichGO`() function from *clusterProfiler* (v4.8.3) package [32], using Nebula DEGs integrated Denisenko samples as the input and queries the Gene Ontology Biological Processes database. The functional enrichment results were visualized with emapplot function from *enrichplot* (v1.20.3) package, with plotting layout set as “kk” [62].

### 4.5 Spot-level deconvolution of cell types

Spatially aware deconvolution was performed for each of the 11 spatially resolved samples using the *CARD* (Conditional AutoregRessive-based Deconvolution, v1.1) method [29].

The reference dataset for *CARD* deconvolution runs was constructed by integrating 5 HGSC 10x genomics single-cell datasets published by Denisenko et al. and 7 additional HGSC 10x genomics scRNAseq datasets generated at HCI. Merged raw count matrices of the single-cell datasets, along with metadata specifying cell type annotations and sample IDs, were used as reference input to *CARD* deconvolution.

### 4.6 HGSC subtype assignment

Subtype assignment to samples was achieved with *consensusOV* (v1.22.0) package using pseudo-bulked expression, generated by summing across raw count matrices across spots within each sample and subsequently library size normalized and log transformed [9]. Molecular subtype classification was attempted using different methods implemented by the package: *consensusOV, Helland, Konecny*, and *Verhaak*.

### 4.7 Cell-cell communication analysis

Cell-cell communication analysis on the Denisenko Visium samples were performed with *COMMOT* (COMMunication analysis by Optimal Transport) [43]. The Visium raw counts were normalized with scanpy with default parameters and log transformed. Initial filtering of potential ligand-receptor pairs was done with *COMMOT* ‘s `filter_lr_database`function, using `Cellchat`as database and `minimum_cell`= 5 as filtering criteria [44]. Cell-cell interaction is inferred with *COMMOT*`spatial_communication`, with interaction distance set as 250.

### 4.8 Cell2location analysis

The *Cell2Location* reference regression model is trained based on the integrated scRNA-seq dataset (see for integration methods). Only cells corresponding to the same donor as the Utah spatial samples from integrated scRNA-seq were retained, and subclustering is applied to Epithelial cell clusters identified in the annotation (**Figure S5**), and then filtered using the gene and cell filtering parameters as suggested by the official documentation (`cell _count_cutoff`= 5, `cell_percentage_cutoff`= 0.03, nonz mean cutoff= 1.12). The estimated expression profile was exported from the trained reference model and passed to the `Cell2location`function with each Visium sample, using `N_cells_per_location`= 5 and `detection_alpha`= 200. The 5th percentile estimate for cell-type abundance computed by *Cell2location* in each spot was plotted.

## 5 Back Matter

### Data Availability

All data and code used in this study are publicly available. The spatial transcriptomics data and single-cell RNA sequencing data generated by HCI can be accessed on dbGaP with study accession id phs002262.v3.p1 (https://www.ncbi.nlm.nih.gov/projects/gap/cgi-bin/study.cgi?study_id=phs002262.v3.p1).

The cellranger/spaceranger processed count matrices and corresponding metadata of the 8 spatial transcriptomics data and 5 single-cell RNA sequencing data published by Denisenko et al. [18] were retrieved from GEO accession number GSE211956 [18].

### Code Availability

Code related code to all analysis used for this manuscript is available on GitHub at https://github.com/wli51/Inter_patient_heterogeneity_HGSOC_spatial_codes. This repository includes all R and Python scripts necessary to replicate the analysis and results presented in this manuscript.

## Acknowledgments

The authors would like to thank several individuals for their assistance in this project. Thanks to the lab members in Kasper Hansen and Stephanie Hicks’s labs for their feedback. We also thank the maintainers of the Joint High Performance Computing Exchange (JHPCE) compute cluster at Johns Hopkins Bloomberg School of Public Health for providing essential computing resources.

Huntsman Cancer Institute, University of Utah: Research reported in this publication utilized the Biorepository and Molecular Pathology Shared Resource, the High-Throughput Genomics and Cancer Bioinformatics Shared Resource, and the Research Informatics Shared Resource at Huntsman Cancer Institute at the University of Utah and was supported by the National Cancer Institute of the National Institutes of Health under Award Number P30CA042014. The content is solely the responsibility of the authors and does not necessarily represent the official views of the NIH.

10x Genomics: We thank the 10x Genomics Visium Spatial Gene Expression Grant Program for the spatial kit and reagents.

Cedars-Sinai: We thank Applied Genomics, Computation & Translational Core at Cedars-Sinai for performing the 10x spatial transcriptomics experiments.

## Author Contributions Statement

- Conceptualization: WL, JAD, CSG, SCH
- Methodology: WL
- Formal Analysis: WL, SCH
- Investigation: WL, SCH
- Resources: LG, JG, JAD
- Data Curation: LG, JAD
- Writing – original draft: WL, SCH
- Writing – review & editing: all authors
- Visualization: WL
- Supervision: JAD, CSG, SCH
- Funding acquisition: JAD, CSG, SCH
- Project Administration: SCH

