## Supplemental Materials for "Characterizing intra- and inter-tumor heterogeneity in Ovarian high-grade serous carcinoma subtypes using single-cell and spatial transcriptomics"

### Contents

1. **Supplementary Tables S1-S4**
2. **Supplementary Figures S1-S12**

| QC-summary | | | | | |
|---|---|---|---|---|---|
| Sample | pre-QC n-cells | post-QC n-cells | pre-QC n-genes | post-QC n-genes | source |
| 16030X2 | 7123 | 3469 | 36601 | 22537 | HCI at Utah |
| 16030X3 | 1533 | 1102 | 36601 | 19931 | HCI at Utah |
| 16030X4 | 6546 | 3260 | 36601 | 23782 | HCI at Utah |
| 19459X1 | 4727 | 3272 | 36601 | 27498 | HCI at Utah |
| 19595X1 | 2595 | 1151 | 36601 | 26289 | HCI at Utah |
| 19833X1 | 2916 | 2536 | 36601 | 25857 | HCI at Utah |
| 19833X2 | 3402 | 2788 | 36601 | 23917 | HCI at Utah |

**Supplementary Table S1: Summary of from Utah HCI scRNA-Seq dataset before and after quality control (QC) filtering.**

| Sample ID | Source | Seuqencing Method | cOV Molecular Subtype |
|---|---|---|---|
| 2B | HCI at Utah | 10x Visium | DIF |
| 3C | HCI at Utah | 10x Visium | DIF |
| 4D | HCI at Utah | 10x Visium | DIF |
| SP1 | Denisenko et. al | 10x Visium | DIF |
| SP2 | Denisenko et. al | 10x Visium | MES |
| SP3 | Denisenko et. al | 10x Visium | MES |
| SP4 | Denisenko et. al | 10x Visium | DIF |
| SP5 | Denisenko et. al | 10x Visium | IMR |
| SP6 | Denisenko et. al | 10x Visium | IMR |
| SP7 | Denisenko et. al | 10x Visium | DIF |
| SP8 | Denisenko et. al | 10x Visium | MES |

**Supplementary Table S2: Sample-level summary of the of the HGSC samples.** For each 10x Genomics Visium capturea area (rows), the table includes the Sample ID, the source or where the tissue was obtained from, the experimental protocol used, and the predicted cOV molecular subtype.

| | | SP1 | | | SP4 | | | SP7 | | |
|---|---|---|---|---|---|---|---|---|---|---|
| Pathway | Ligand-Receptor | mean | 1st quartile | 3rd quartile | mean | 1st quartile | 3rd quartile | mean | 1st quartile | 3rd quartile |
| MK | MDK.ITGA6_ITGB1 | 2.76 | 0.03 | 4.22 | 1.21 | 0.00 | 0.25 | 1.26 | 0.00 | 0.13 |
| | MDK.LRP1 | 9.89 | 3.10 | 13.66 | 8.37 | 0.30 | 13.06 | 8.39 | 0.00 | 11.54 |
| | MDK.NCL | 8.10 | 1.48 | 10.98 | 15.56 | 6.40 | 22.11 | 11.23 | 0.16 | 15.64 |
| | MDK.SDC1 | 7.50 | 0.69 | 10.66 | 2.52 | 0.00 | 2.56 | 2.09 | 0.00 | 0.94 |
| | MDK.SDC2 | 2.62 | 0.06 | 3.35 | 3.73 | 0.00 | 5.66 | 3.01 | 0.00 | 2.92 |
| | MDK.SDC4 | 3.67 | 0.08 | 5.23 | 6.21 | 0.12 | 10.16 | 11.86 | 0.11 | 16.06 |
| MIF | MIF.ACKR3 | 2.54 | 0.00 | 3.57 | 0.76 | 0.00 | 0.00 | 0.94 | 0.00 | 0.00 |
| | MIF.CD74_CD44 | 1.70 | 0.00 | 1.63 | 2.82 | 0.00 | 1.90 | 6.73 | 0.00 | 9.46 |
| | MIF.CD74_CXCR4 | 5.08 | 0.00 | 7.93 | 5.03 | 0.00 | 5.81 | 0.80 | 0.00 | 0.00 |
| SPP1 | SPP1.CD44 | 1.46 | 0.00 | 0.21 | 0.79 | 0.00 | 0.00 | 8.23 | 0.00 | 9.94 |
| | SPP1.ITGA8_ITGB1 | 0.03 | 0.00 | 0.00 | 0.01 | 0.00 | 0.00 | 0.13 | 0.00 | 0.00 |
| | SPP1.ITGA9_ITGB1 | 0.65 | 0.00 | 0.00 | 0.35 | 0.00 | 0.00 | 0.20 | 0.00 | 0.00 |
| | SPP1.ITGAV_ITGB1 | 2.00 | 0.00 | 2.08 | 0.66 | 0.00 | 0.00 | 1.57 | 0.00 | 0.35 |
| | SPP1.ITGAV_ITGB5 | 0.85 | 0.00 | 0.01 | 0.43 | 0.00 | 0.00 | 1.65 | 0.00 | 0.24 |
| | SPP1.ITGAV_ITGB6 | 0.08 | 0.00 | 0.00 | 0.06 | 0.00 | 0.00 | 0.73 | 0.00 | 0.00 |
| SEMA3 | SEMA3A.NRP1_PLXNA1 | 0.16 | 0.00 | 0.00 | 0.05 | 0.00 | 0.00 | 0.21 | 0.00 | 0.00 |
| | SEMA3A.NRP1_PLXNA2 | 0.08 | 0.00 | 0.00 | 0.04 | 0.00 | 0.00 | 0.10 | 0.00 | 0.00 |
| | SEMA3A.NRP1_PLXNA4 | 0.06 | 0.00 | 0.00 | 0.01 | 0.00 | 0.00 | 0.01 | 0.00 | 0.00 |
| | SEMA3B.NRP1_PLXNA1 | 0.39 | 0.00 | 0.00 | 0.14 | 0.00 | 0.00 | 0.36 | 0.00 | 0.00 |
| | SEMA3B.NRP1_PLXNA2 | 0.21 | 0.00 | 0.00 | 0.13 | 0.00 | 0.00 | 0.13 | 0.00 | 0.00 |
| | SEMA3B.NRP1_PLXNA4 | 0.13 | 0.00 | 0.00 | 0.03 | 0.00 | 0.00 | 0.02 | 0.00 | 0.00 |
| | SEMA3B.NRP2_PLXNA1 | 0.57 | 0.00 | 0.01 | 0.96 | 0.00 | 0.00 | 0.86 | 0.00 | 0.00 |
| | SEMA3B.NRP2_PLXNA2 | 0.23 | 0.00 | 0.00 | 0.57 | 0.00 | 0.00 | 0.30 | 0.00 | 0.00 |
| | SEMA3B.NRP2_PLXNA4 | 0.23 | 0.00 | 0.00 | 0.06 | 0.00 | 0.00 | 0.03 | 0.00 | 0.00 |
| | SEMA3C.NRP1_NRP2_PLXND1 | 0.17 | 0.00 | 0.00 | 0.10 | 0.00 | 0.00 | 0.28 | 0.00 | 0.00 |
| | SEMA3C.NRP1_PLXNA1 | 0.20 | 0.00 | 0.00 | 0.21 | 0.00 | 0.00 | 0.24 | 0.00 | 0.00 |
| | SEMA3C.NRP1_PLXNA2 | 0.09 | 0.00 | 0.00 | 0.13 | 0.00 | 0.00 | 0.12 | 0.00 | 0.00 |
| | SEMA3C.NRP1_PLXNA4 | 0.06 | 0.00 | 0.00 | 0.03 | 0.00 | 0.00 | 0.01 | 0.00 | 0.00 |
| | SEMA3C.NRP2_PLXNA1 | 0.20 | 0.00 | 0.00 | 0.63 | 0.00 | 0.00 | 0.61 | 0.00 | 0.00 |
| | SEMA3C.NRP2_PLXNA2 | 0.09 | 0.00 | 0.00 | 0.43 | 0.00 | 0.00 | 0.27 | 0.00 | 0.00 |
| | SEMA3C.NRP2_PLXNA4 | 0.10 | 0.00 | 0.00 | 0.08 | 0.00 | 0.00 | 0.03 | 0.00 | 0.00 |
| | SEMA3C.PLXND1 | 1.55 | 0.00 | 0.00 | 2.15 | 0.00 | 0.21 | 3.11 | 0.00 | 3.00 |
| | SEMA3D.NRP1_PLXNA1 | 0.03 | 0.00 | 0.00 | 0.01 | 0.00 | 0.00 | 0.00 | 0.00 | 0.00 |
| | SEMA3D.NRP1_PLXNA2 | 0.03 | 0.00 | 0.00 | 0.01 | 0.00 | 0.00 | 0.00 | 0.00 | 0.00 |
| | SEMA3D.NRP1_PLXNA4 | 0.01 | 0.00 | 0.00 | 0.00 | 0.00 | 0.00 | 0.00 | 0.00 | 0.00 |
| | SEMA3D.NRP2_PLXNA1 | 0.04 | 0.00 | 0.00 | 0.02 | 0.00 | 0.00 | 0.01 | 0.00 | 0.00 |
| | SEMA3D.NRP2_PLXNA2 | 0.02 | 0.00 | 0.00 | 0.00 | 0.00 | 0.00 | 0.00 | 0.00 | 0.00 |
| | SEMA3D.NRP2_PLXNA4 | 0.02 | 0.00 | 0.00 | 0.00 | 0.00 | 0.00 | 0.00 | 0.00 | 0.00 |
| | SEMA3G.NRP2_PLXNA1 | 0.26 | 0.00 | 0.00 | 0.05 | 0.00 | 0.00 | 0.04 | 0.00 | 0.00 |
| | SEMA3G.NRP2_PLXNA2 | 0.11 | 0.00 | 0.00 | 0.04 | 0.00 | 0.00 | 0.02 | 0.00 | 0.00 |
| | SEMA3G.NRP2_PLXNA4 | 0.13 | 0.00 | 0.00 | 0.01 | 0.00 | 0.00 | 0.01 | 0.00 | 0.00 |
| ncWNT | WNT11.FZD1 | 0.28 | 0.00 | 0.00 | 0.11 | 0.00 | 0.00 | 0.20 | 0.00 | 0.00 |
| | WNT11.FZD2 | 0.32 | 0.00 | 0.00 | 0.09 | 0.00 | 0.00 | 0.10 | 0.00 | 0.00 |
| | WNT11.FZD3 | 0.35 | 0.00 | 0.00 | 0.14 | 0.00 | 0.00 | 0.14 | 0.00 | 0.00 |
| | WNT11.FZD4 | 0.24 | 0.00 | 0.00 | 0.11 | 0.00 | 0.00 | 0.19 | 0.00 | 0.00 |
| | WNT11.FZD7 | 0.42 | 0.00 | 0.00 | 0.12 | 0.00 | 0.00 | 0.11 | 0.00 | 0.00 |
| | WNT11.FZD8 | 0.14 | 0.00 | 0.00 | 0.01 | 0.00 | 0.00 | 0.06 | 0.00 | 0.00 |
| | WNT5A.FZD1 | 0.46 | 0.00 | 0.00 | 1.08 | 0.00 | 0.00 | 0.37 | 0.00 | 0.00 |
| | WNT5A.FZD2 | 0.58 | 0.00 | 0.00 | 1.26 | 0.00 | 0.00 | 0.25 | 0.00 | 0.00 |
| | WNT5A.FZD3 | 0.60 | 0.00 | 0.00 | 1.93 | 0.00 | 0.11 | 0.37 | 0.00 | 0.00 |
| | WNT5A.FZD4 | 0.38 | 0.00 | 0.00 | 1.45 | 0.00 | 0.00 | 0.41 | 0.00 | 0.00 |
| | WNT5A.FZD7 | 0.77 | 0.00 | 0.00 | 1.20 | 0.00 | 0.00 | 0.27 | 0.00 | 0.00 |
| | WNT5A.FZD8 | 0.25 | 0.00 | 0.00 | 0.09 | 0.00 | 0.00 | 0.15 | 0.00 | 0.00 |
| | WNT5A.MCAM | 0.93 | 0.00 | 0.00 | 0.53 | 0.00 | 0.00 | 0.30 | 0.00 | 0.00 |
| PTN | PTN.NCL | 5.74 | 0.00 | 8.25 | 0.40 | 0.00 | 0.00 | 0.20 | 0.00 | 0.00 |
| | PTN.SDC1 | 4.47 | 0.00 | 6.98 | 0.16 | 0.00 | 0.00 | 0.08 | 0.00 | 0.00 |
| | PTN.SDC2 | 1.71 | 0.00 | 2.35 | 0.17 | 0.00 | 0.00 | 0.08 | 0.00 | 0.00 |
| | PTN.SDC3 | 1.53 | 0.00 | 1.52 | 0.14 | 0.00 | 0.00 | 0.09 | 0.00 | 0.00 |
| | PTN.SDC4 | 2.32 | 0.00 | 3.33 | 0.25 | 0.00 | 0.00 | 0.19 | 0.00 | 0.00 |
| PDGF | PDGFA.PDGFRA | 0.18 | 0.00 | 0.00 | 0.62 | 0.00 | 0.00 | 0.40 | 0.00 | 0.00 |
| | PDGFA.PDGFRB | 0.38 | 0.00 | 0.00 | 0.39 | 0.00 | 0.00 | 0.52 | 0.00 | 0.00 |
| | PDGFC.PDGFRA | 1.30 | 0.00 | 0.00 | 3.70 | 0.00 | 1.23 | 0.81 | 0.00 | 0.00 |
| | PDGFD.PDGFRB | 0.48 | 0.00 | 0.00 | 0.37 | 0.00 | 0.00 | 0.22 | 0.00 | 0.00 |
| COMPLEMENT | C3.C3AR1 | 0.57 | 0.00 | 0.00 | 0.57 | 0.00 | 0.00 | 0.78 | 0.00 | 0.00 |
| | C3.ITGAM_ITGB2 | 0.07 | 0.00 | 0.00 | 0.05 | 0.00 | 0.00 | 0.31 | 0.00 | 0.00 |
| | C3.ITGAX_ITGB2 | 0.10 | 0.00 | 0.00 | 0.02 | 0.00 | 0.00 | 0.49 | 0.00 | 0.00 |
| CXCL | CXCL10.ACKR1 | 0.44 | 0.00 | 0.00 | 0.01 | 0.00 | 0.00 | 0.10 | 0.00 | 0.00 |
| | CXCL12.ACKR3 | 0.45 | 0.00 | 0.00 | 0.22 | 0.00 | 0.00 | 0.27 | 0.00 | 0.00 |
| | CXCL12.CXCR4 | 1.35 | 0.00 | 0.00 | 0.84 | 0.00 | 0.00 | 0.33 | 0.00 | 0.00 |
| | CXCL2.ACKR1 | 0.07 | 0.00 | 0.00 | 0.13 | 0.00 | 0.00 | 0.22 | 0.00 | 0.00 |
| | CXCL8.ACKR1 | 0.01 | 0.00 | 0.00 | 0.18 | 0.00 | 0.00 | 0.47 | 0.00 | 0.00 |

**Supplementary Table S3: Summary table for ligand-receptor outgoing signal strength from top (active) signaling pathways for DIF Visium samples (SP1, SP4, SP7), organized by Pathway**. Values represent the mean, first quartile, and third quartile communication scores for each ligand–receptor pair within the indicated pathway, as inferred from cell–COMMOT [44] cell communication analysis. Higher values (length of highlight lenghts) indicate stronger predicted signaling activity from the sender cell population toward all potential receiver populations in the dataset.

| QC-summary | | | | | |
|---|---|---|---|---|---|
| Sample | pre-QC n-cells | post-QC n-cells | pre-QC n-genes | post-QC n-genes | source |
| Y2 | 328 | 328 | 19915 | 13377 | Denisenko et. al |
| Y3 | 6384 | 6384 | 19915 | 17247 | Denisenko et. al |
| Y5 | 7588 | 7588 | 19915 | 17618 | Denisenko et. al |
| MJ10 | 426 | 426 | 19915 | 15248 | Denisenko et. al |
| MJ11 | 1956 | 1956 | 19915 | 17591 | Denisenko et. al |

**Supplementary Table S4: Summary of from Denisenko et al. scRNA-Seq dataset before and after quality control (QC) filtering.**

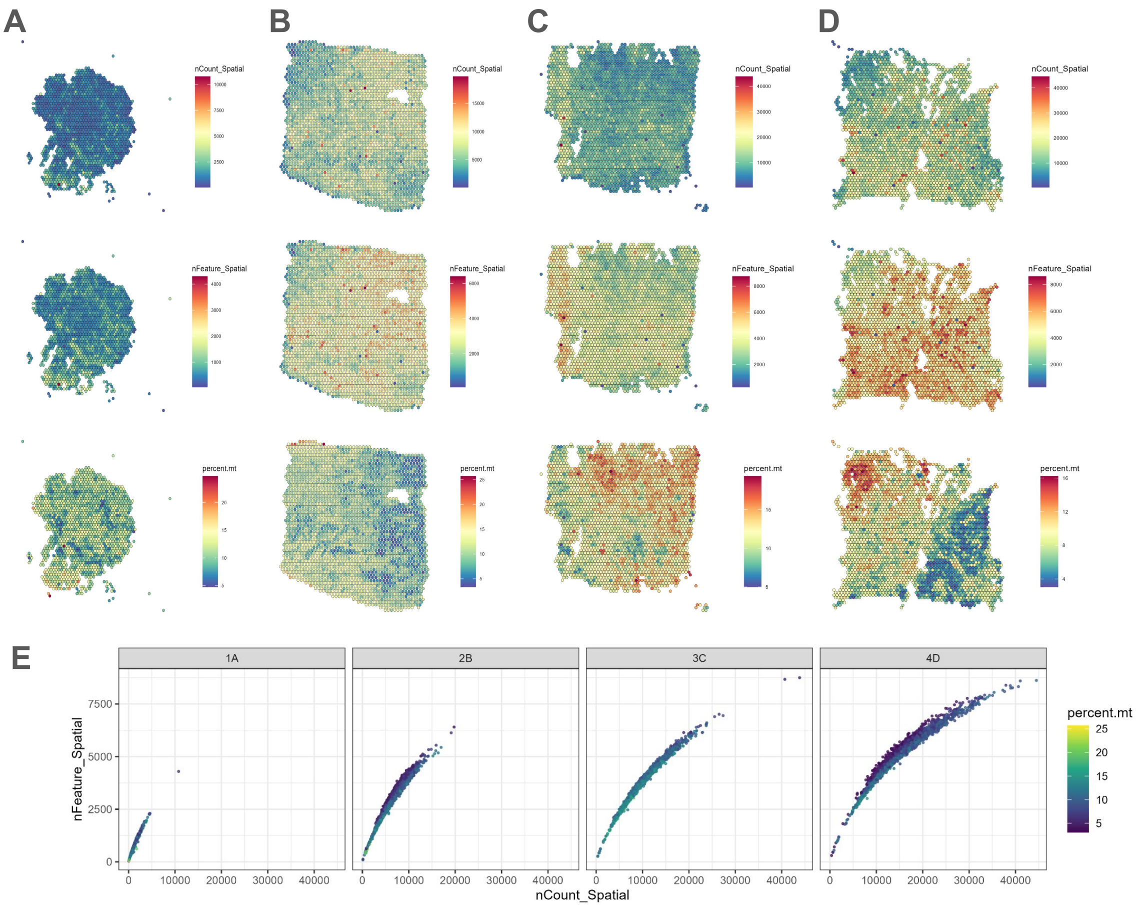


**Supplementary Figure S1: Quality control metrics for $N$=4 tissue sections.** Distribution of total UMIs per capture area for tumor sample 16030X2. Capture area 1A was removed from downstream analysis due to low total number of UMI. **(A-D)** Spatial visualization of spot-wise UMI count (top), expressed gene count (middle) and mitochondrial gene concentration (bottom) across the four capture areas **(E)** Summary scatter plot for the same quality control statistic across all capture areas

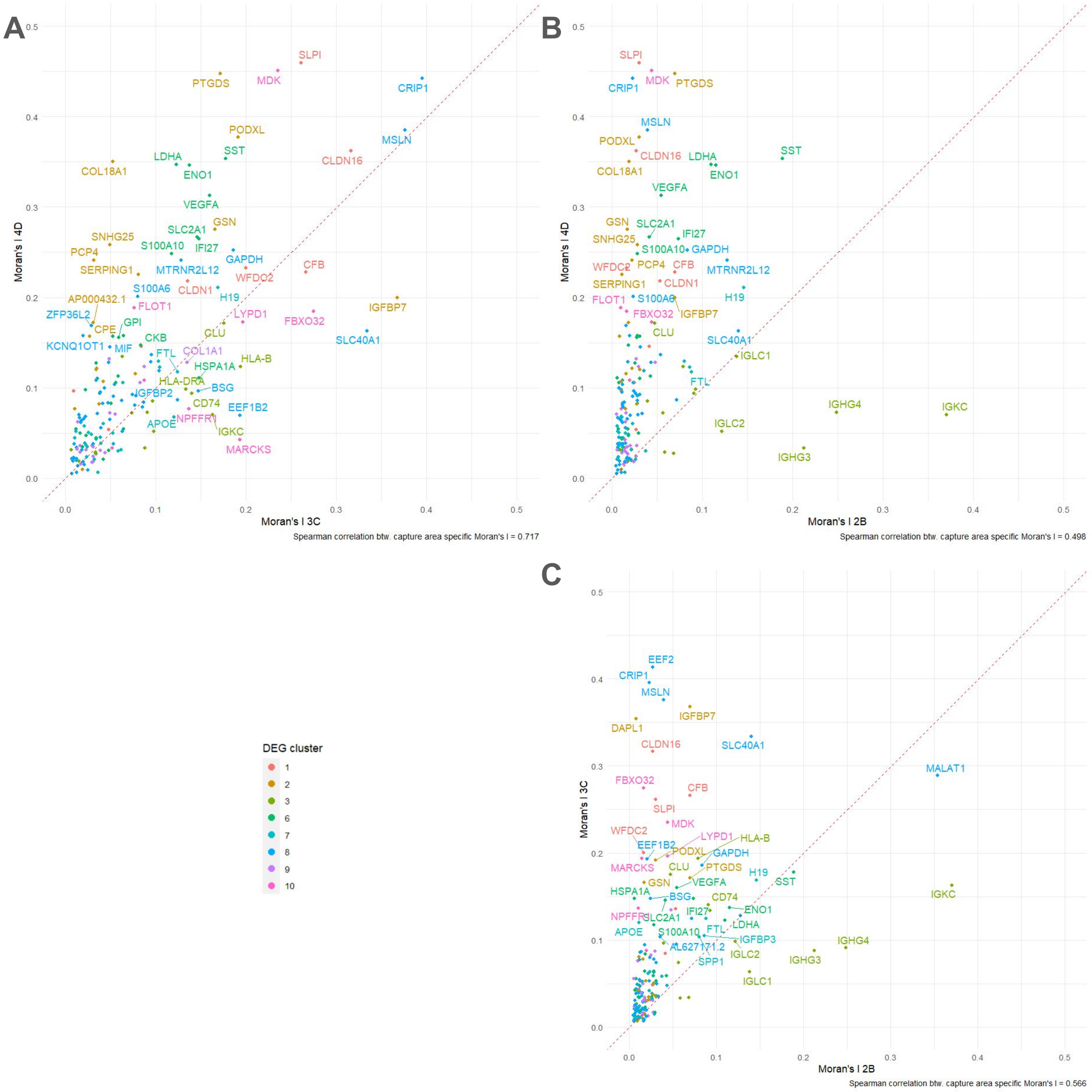


**Supplementary Figure S2: HGSC identify different spatially variable genes.** (**A-C**) Pairwise scatterplots between pairs of $N$=3 capture areas with the spatially variable genes detected using Moran's I statistic, gene name labels are shown only for genes with absolute Moran's I exceeding 0.1 in either sample. The genes names and scatter points were colored in accordance to the NMF cluster from which the gene is identified as DEG.

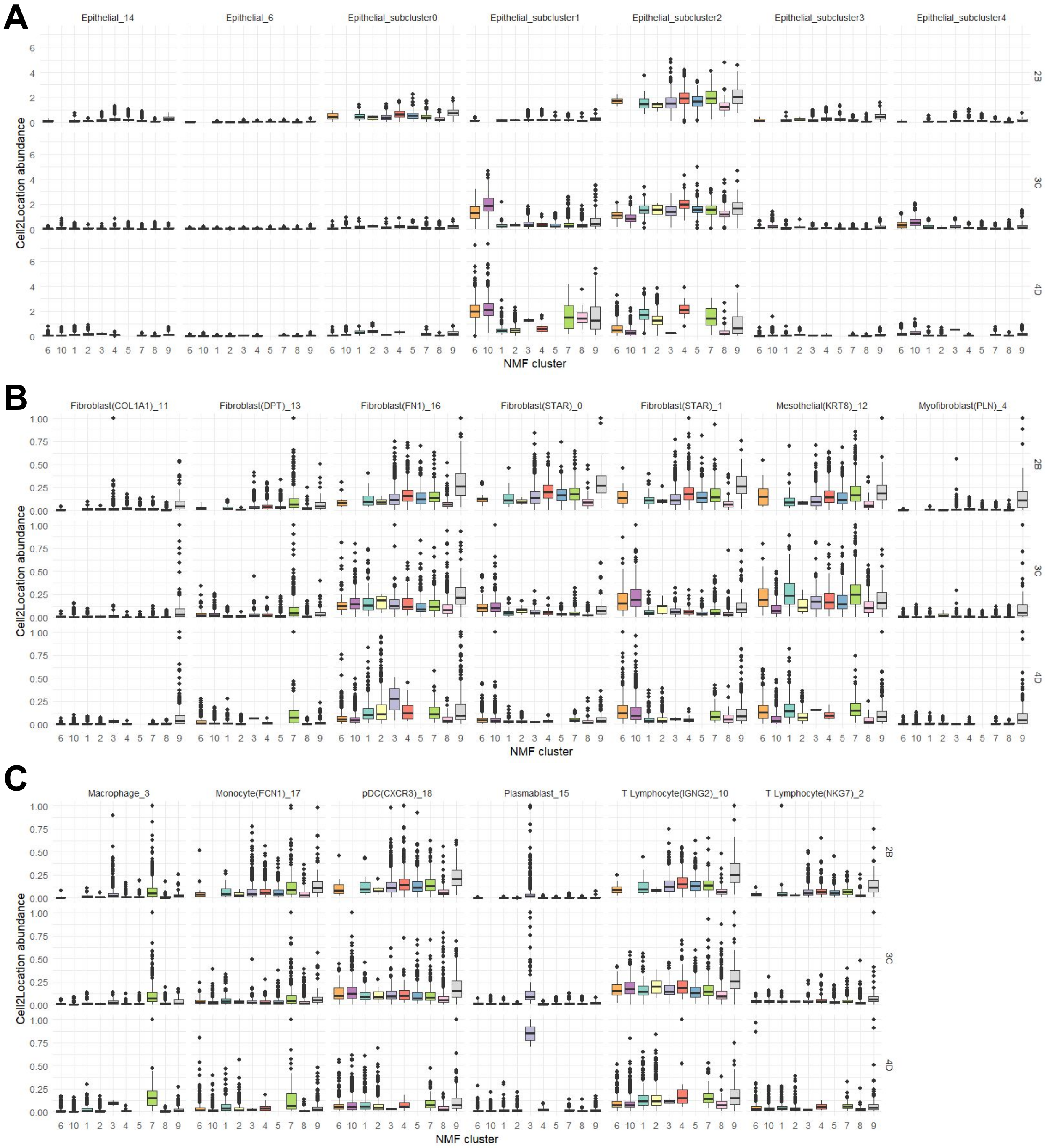


**Supplementary Figure S3:** (Caption next page.)

**Supplementary Figure S3: Cell2location epithelial cell abundance.** Summary statistics of 5th percentile cell-type abundance predictions by *cell2location* mapping of clustered single cell data for **(A)** 2B, **(B)** 3C and **(C)** 4D. The spatial-domain formed by high-abundance Epithelial-subcluster1 subpopulations in 3C and 4D corresponds well with the NMF clustering scheme (cluster 6 and 10) from Figure 1. Fibroblast(STAR) populations also seemed to be exclusively abundant in clusters 6 and 10 while higher Fibroblast(FN1) proportions are associated with other NMF clusters.

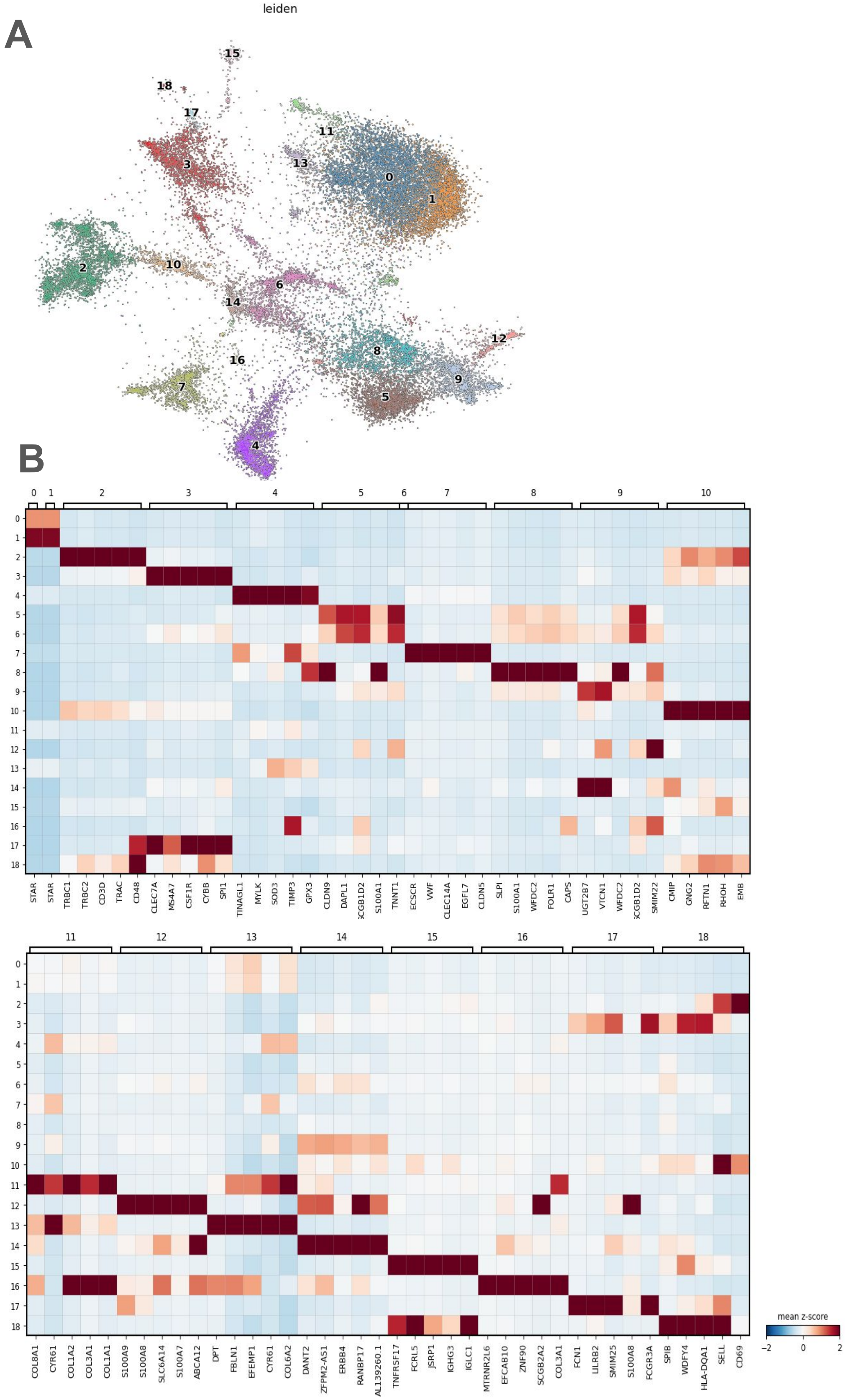


**Supplementary Figure S4:** (Caption next page.)

**Supplementary Figure S4: Annotated single cell dataset summary. (A)** UMAP projection of annotated integrated single cell dataset that is used as reference in cell2location mapping and deconvolution. **(B)** Top 5 DEGs identified for each cluster thruough the bayesian DE function of scVI, in descending ordered of Bayes Factor.

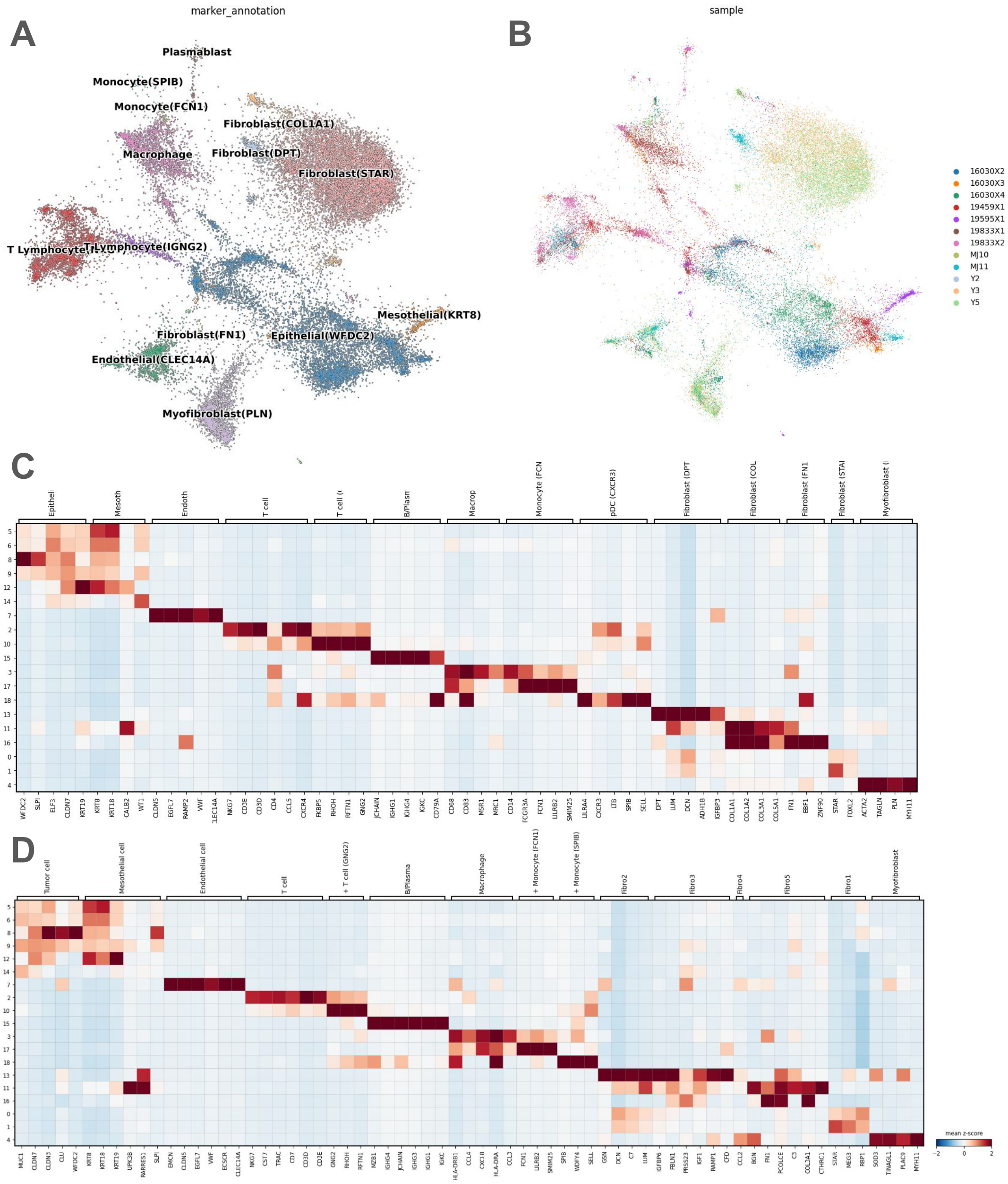


**Supplementary Figure S5: Annotated scRNA-seq data from the integrated Utah samples and Denisenko et al. [18] samples.** **(A-B)** UMAP visualization of the manual annotation and origin of integrated cells, used as reference for deconvolution. The integrated scRNA-seq dataset includes 7 Utah and 5 Denisenko et al. [18] HGSC samples. Integration is carried out with scVI. **(C)** Dotplot illustration of marker genes corresponding to each of the annotated cell types. **(D)** Dotplot illustration of the markers corresponding to the cell type annotation reported by Denisenko et. al. Note that the scVI integration and annotation differed from the Denisenko et al. [18] reference annotation primarily in fibroblast subpopulations. The Denisenko et al. [18] Fibro5 (*FN1*, *COL3A1*) seemed to best correspond to the scVI Fibroblast (*COL1A1*), and Denisenko et al. [18] Fibro2 (*RBP1*, *DCN*) best corresponded scVI Fibroblast (*DPT*), Alistair Fibro3 (*RAMP1*, *CFD*) best corresponded to scVI Fibroblast (*RAMP1*), and Alistair Myofibroblast corresponded to scVI Myofibroblast (*PLN*; *ACTA2*). There seemed to be a lack of correspondence between Alistair Fibro1 and Fibro4 with any of the scVI fibroblast types.

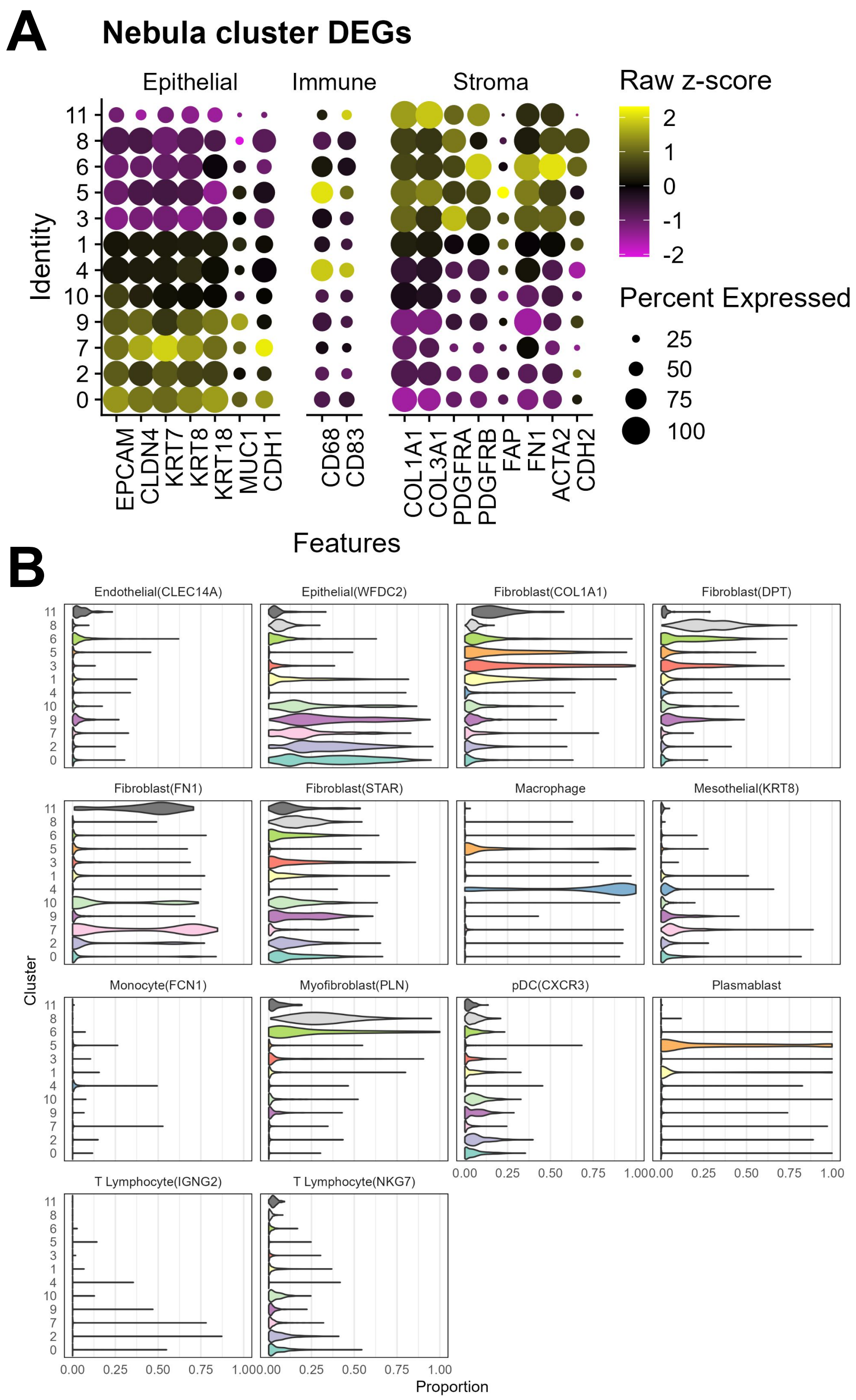


**Supplementary Figure S6: Canonical epithelial vs mesenchymal marker expression in the integrated dataset. (A)** Dot plot showing expression of epithelial and mesenchymal markers. We found that clusters 0, 2, 7, 9 and 10 expressed higher epithelial markers and clusters 3, 5, 6, 8, 11 express higher mesenchymal markers.**(B)** Violin plot showing deconvolution proportions comparing epitheliat-dominant clusters (0, 2, 7, 9, 10) with fibroblast-dominant clusters (3, 5, 6, 8, 11).

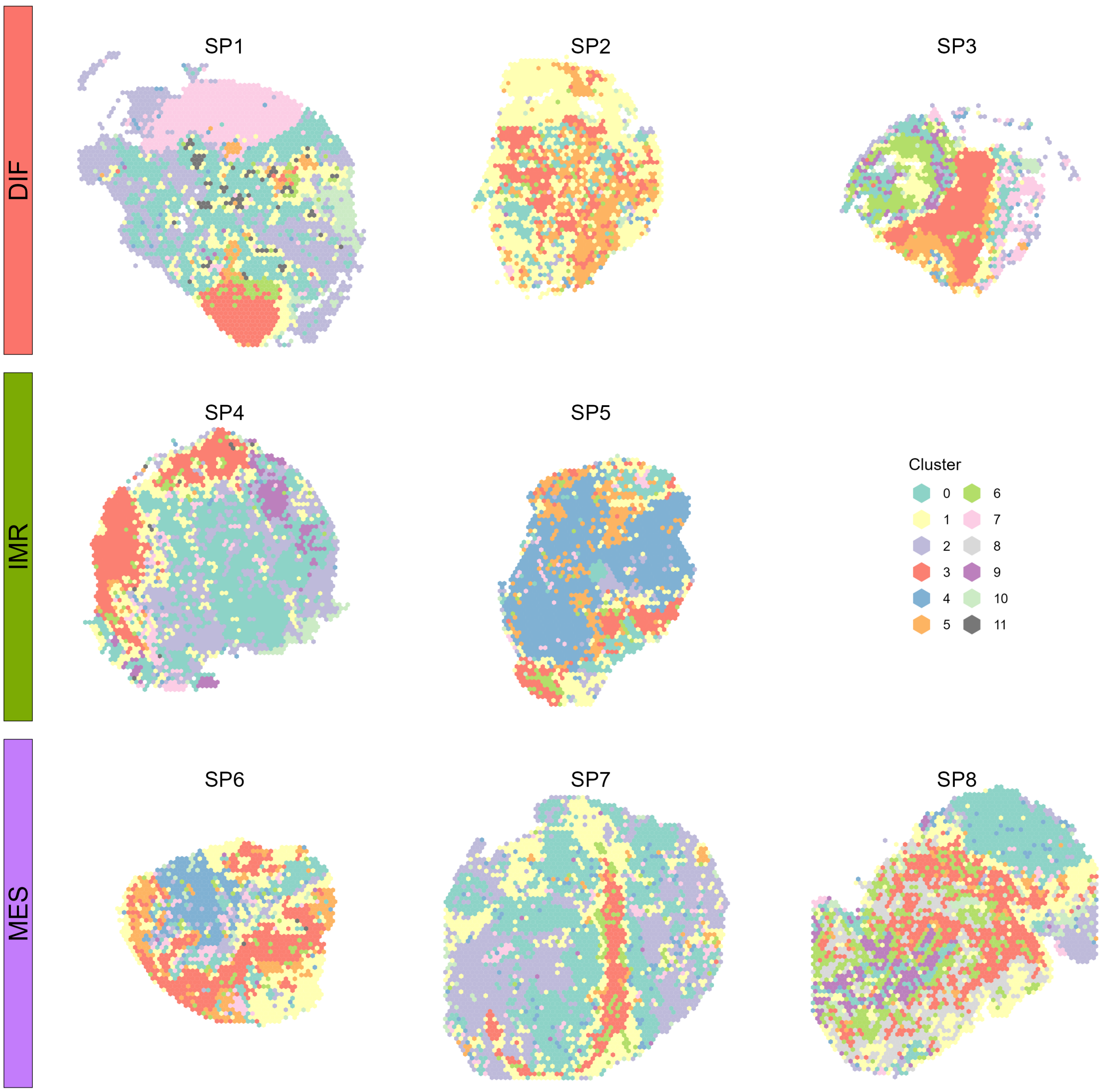


**Supplementary Figure S7: Integrated clustering in space** Clustering scheme in space for the Denisenko et al. [18] integration. In total 12 clusters are identified. Notably, cluster 1 (yellow) occurred almost exclusively between epithelial-dominant (teal, pink, blue-gray) and fibroblast-dominant clusters (red, orange, green, light gray).

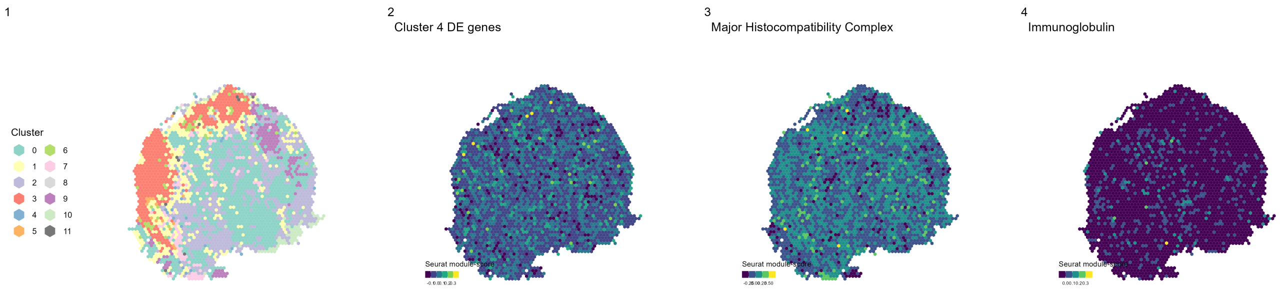


**Supplementary Figure S8: Denisenko et al. Sample SP4 (DIF) Spatial Domain** Immunoglobulin and major histocompatibility complex genes scores visualization for SP4, which is not included in **Figures** 4. Compared to the other DIF samples, SP4 displays less co-expression of MHC and IG modules in space.

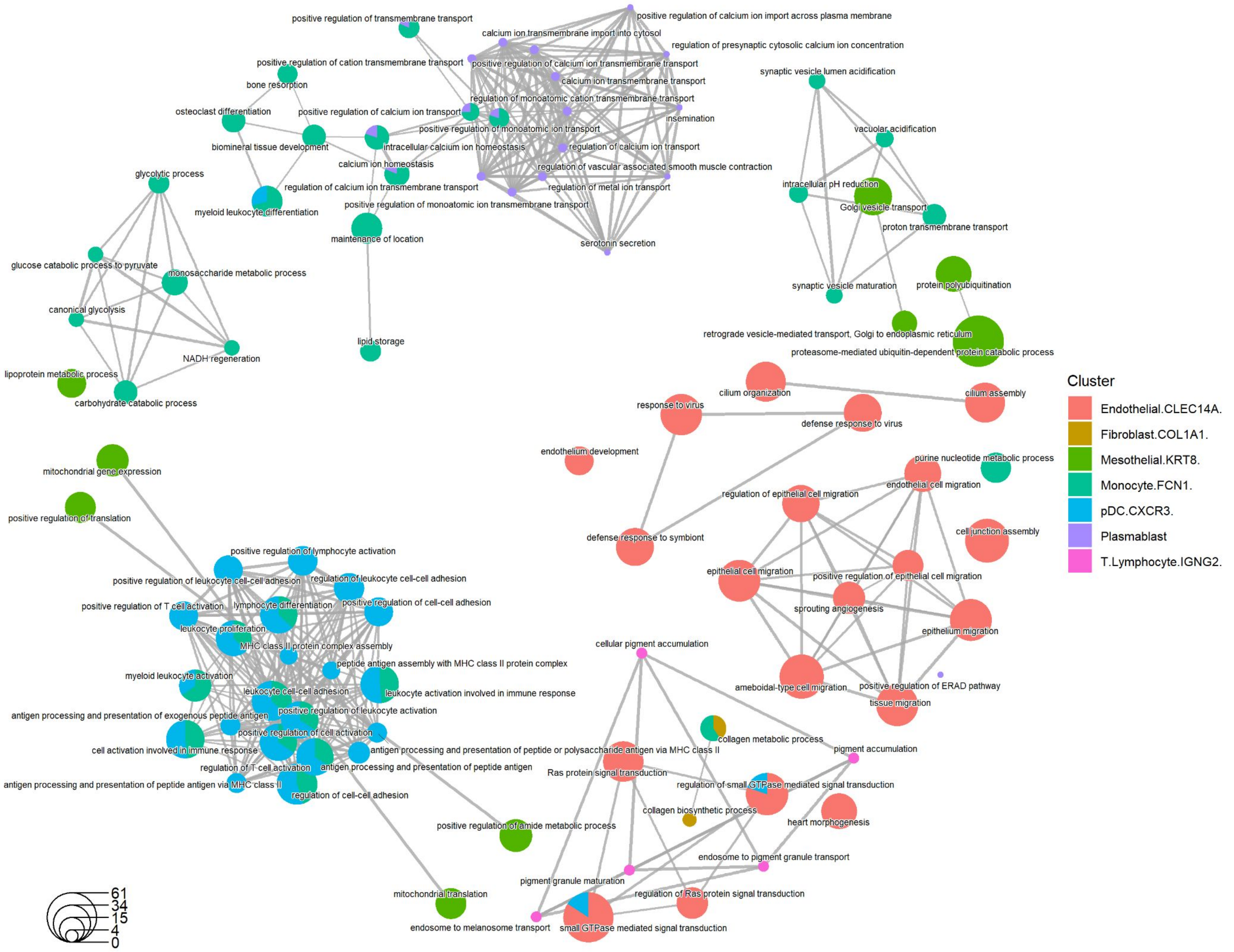


**Supplementary Figure S9:** Gene concept network plot showing the GO (biological pathway) results based on genes whose expression is positively associated with the estimated proportion of cell types from spot-level deconvolution, colored by cell types.

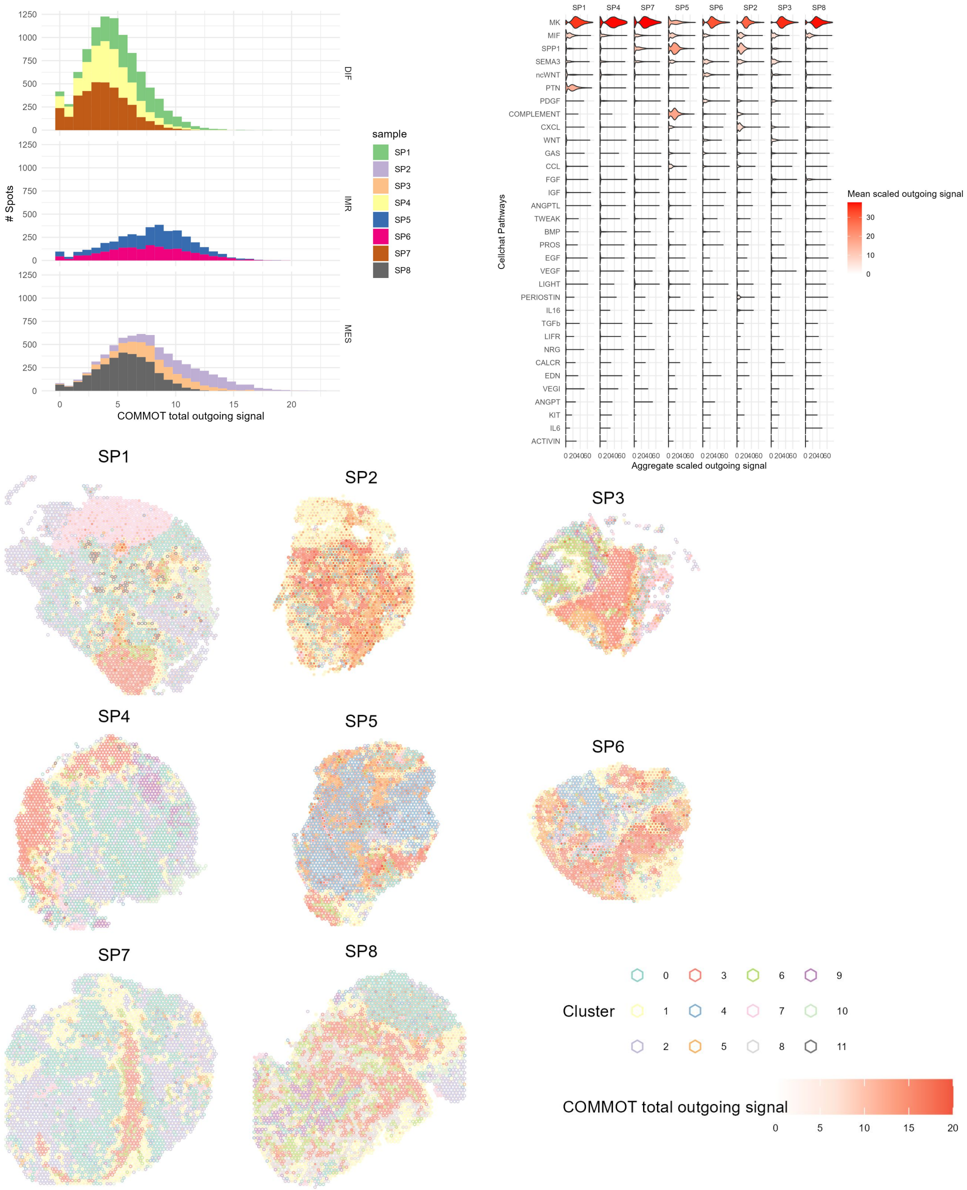

**Supplementary Figure S10: COMMOT ligand-receptor total signaling summary.** **(A)** Histogram indicating distribution of spot total outgoing signals in the $N$=8 Denisenko et al. [18] Visium samples, grouped by molecular subtype. The magnitude and distribution of total outgoing signal is remarkably similar between samples of the same subtype. In general, the IMR samples are more active compared to the MES and DIF samples. **(B)** Violin plot summarizing the distribution of scaled outgoing signaling in each pathway across all samples. **(C)** Total signaling activity in space. Signaling activity is represented with fill color, while cluster is represented as boundary line of spots. A substantial amount of signaling occurs near the boundary of clusters.

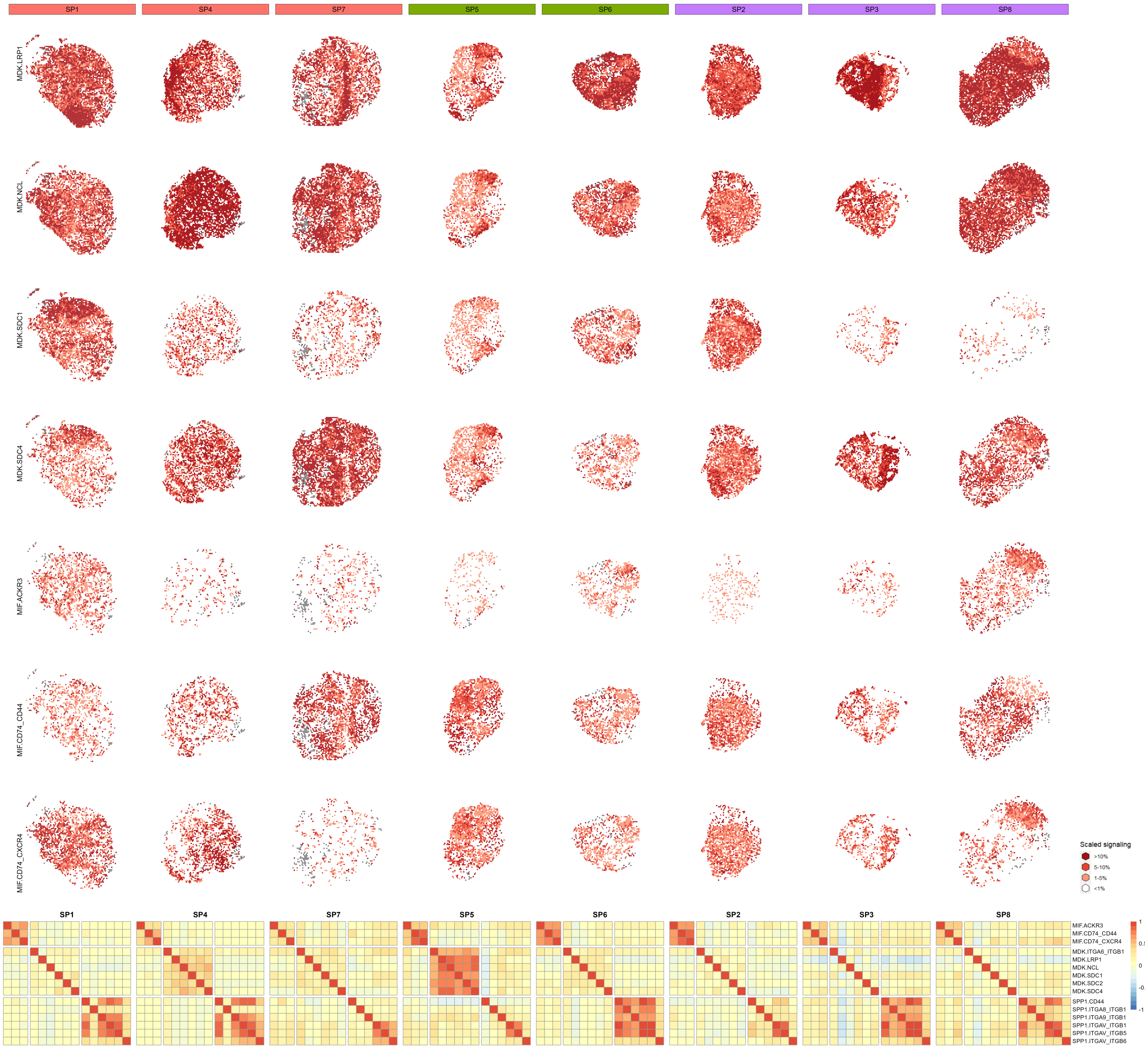


**Supplementary Figure S11: COMMOT ligand-receptor signaling correlation.** [Spatial plots, top] Scaled signaling activities in select ligand receptor pairs. Each sample is described by one column, indicated by the column label. Each selected ligand receptor pair is represented by the same row. [Heatmap, bottom] Spearman correlation of ligand receptor pairs from the top 3 most active pathways (MIF, MK, SPP1). The correlation structure appeared to be highly sample specific and blacked subtype specific patterns, except for the MES specific anti-correlation between MDK-LRP1 signaling with other MK, MIF and SPP1 pathway ligand receptor pairs (heatmap SP3, SP8). Notably, ligand receptor activities from the MK pathway clearly colocalize to different cluster/spatial domains of the tumor tissue in SP1, SP7, SP3 and SP8, corresponding to the lack of inter-pathway correlation in the corresponding heatmaps.

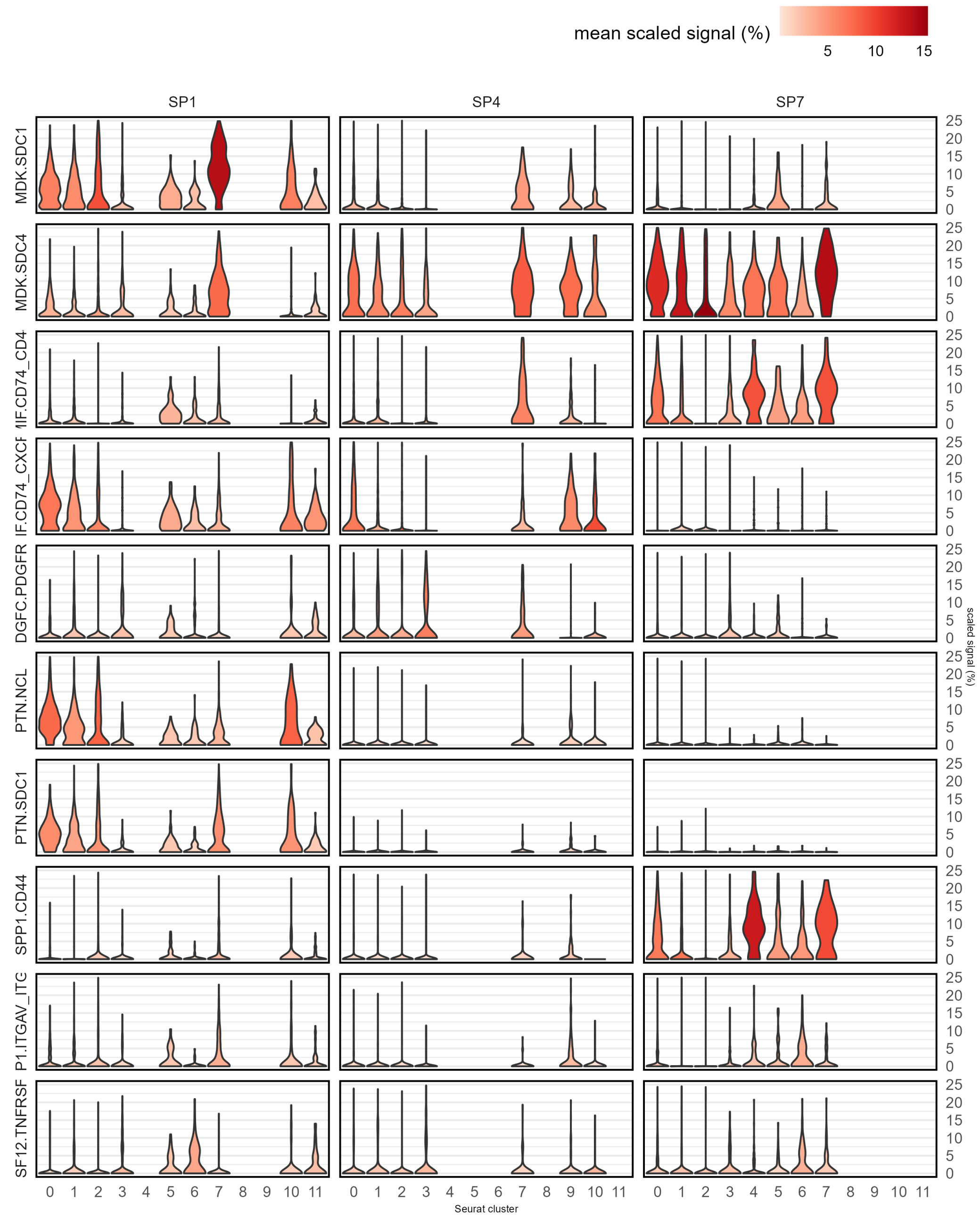


**Supplementary Figure S12: COMMOT select ligand-receptor pair signaling strength summary for DIF samples** The violin plot represents the distribution of scaled outgoing signal strength. The fill of each violin corresponds to the mean scaled outgoing signal strength in spatial domain of sample. Domains with fewer than 50 spots per sample omitted.
